# Parallel connectivity and genomic divergence but heterogeneous demographic history in sympatric Mediterranean clingfishes

**DOI:** 10.64898/2026.06.16.732606

**Authors:** Maximilian Wagner, Stephan Koblmüller, Ariane Droin, Nadine Klar, Sandra Bračun, Stamatis Zogaris, Kristina M. Sefc, Hannes Svardal

## Abstract

Understanding the extent to which marine connectivity can be predicted from shared oceanographic and biogeographic conditions remains a central challenge in marine ecology and evolution. Comparative analyses of sympatric species provide a powerful framework for testing whether common environmental contexts generate repeatable patterns of population structure and connectivity and if this also unfolds on a genomic level. Here, we investigated this question in the endemic Mediterranean clingfish genus *Gouania*, a group of cryptobenthic gravel-beach specialists with limited adult mobility and a short pelagic larval phase. We combined whole-genome resequencing of two sympatric eastern Mediterranean species (*G. orientalis* and *G. hofrichteri*) around Crete and Kythira with Lagrangian particle drift simulations, mitochondrial phylogeography across all five *Gouania* species and demographic reconstructions using MSMC2. Passive drift simulations predicted strong isolation of Kythira and restricted exchange between northern and southern Crete. These predictions closely matched genome-wide population structure in both species, revealing highly concordant patterns of connectivity despite independent evolutionary histories. Shared population contrasts were furthermore associated with a significant excess of parallel genomic divergence, with both genomic windows and candidate genes overlapping more frequently than expected by chance. Nevertheless, most differentiated genomic regions remained species-specific and functional enrichment analyses identified no dominant shared pathways, suggesting a largely polygenic basis of divergence. In contrast to the strong concordance observed for contemporary connectivity and genomic divergence, mitochondrial phylogeographic patterns and species-level demographic histories were considerably more heterogeneous across the genus. Together, our results demonstrate that the predictability of marine population structure is scale-dependent: shared seascapes can generate repeatable patterns of connectivity and partially parallel genomic divergence, whereas deeper phylogeographic and demographic histories remain strongly influenced by species-specific historical processes.

## INTRODUCTION

Understanding how marine populations are connected is central to guiding conservation planning and designing effective spatial management, particularly in regions undergoing rapid environmental and coastal change such as the Mediterranean Sea (Giakoumi et al., 2013; Jahnke et al., 2017). Yet predicting gene flow in marine systems remains challenging because connectivity is shaped by the interaction between intrinsic species traits and extrinsic seascape or oceanographic processes. In species with limited adult mobility, dispersal often depends on a pelagic larval phase, whose realised contribution to gene flow is influenced by reproductive timing, pelagic larval duration, larval behaviour, self-recruitment, habitat distribution and physical transport by currents (Cowen et al., 2007; Klein et al., 2016; Ollé-Vilanova et al., 2022; Pascual et al., 2017; Sefc et al., 2020). As a result, dispersal ability does not always translate directly into genetic connectivity: some taxa with seemingly limited dispersal potential show high connectivity, whereas others with apparently greater dispersal potential exhibit pronounced population structure (Weersing & Toonen, 2009). In addition, historical demographic processes can leave persistent genomic signatures that complicate the interpretation of contemporary population structure and realised connectivity (Arranz et al., 2021; Hellberg, 2009; Selkoe et al., 2008).

The Mediterranean Sea provides an excellent setting for disentangling the processes shaping marine connectivity because contemporary oceanographic structure and historical biogeographic legacies operate together across multiple spatial and temporal scales. Present-day circulation is highly structured and seasonally dynamic, with regional gyres, jets, fronts and semi-enclosed basins that can generate dispersal barriers, asymmetric connectivity and fine-scale population structure even over short geographic distances (Cowen et al., 2007; Schroeder et al., 2012; Sefc et al., 2020). At the same time, historical events such as the Messinian salinity crisis around five million years ago and Quaternary sea-level fluctuations have shaped Mediterranean biogeographic boundaries, demographic histories and divergence times across marine taxa (Agiadi et al., 2024; Bianchi et al., 2012; Deli et al., 2019; Koblmüller et al., 2015; Kousteni et al., 2015). Thus, Mediterranean marine populations are expected to carry signatures of both contemporary dispersal processes and deeper historical events.

Comparing sympatric species offers a powerful way to test the predictability of marine connectivity. Because such species occupy the same regional seascape, many extrinsic drivers of dispersal and demographic history are shared (Dawson, 2012). Concordant patterns among species would suggest that oceanographic and biogeographic processes strongly shape realised connectivity, whereas discordant patterns would point to the importance of species-specific traits, ecological differences, or demographic stochasticity (Kato et al., 2025). Yet, although this comparative logic is central to phylogeography, it has rarely been applied at the genomic scale to closely related sympatric species. Such systems offer an additional advantage: because orthologous genomic regions can be compared within a shared evolutionary framework, they allow tests not only of whether shared seascapes or oceanography generate similar population structure, but also of whether shared environmental pressures are associated with parallel genomic landscapes of divergence, for example through selection on the same genes or pathways (Wagner et al., 2026).

The endemic Mediterranean clingfish genus *Gouania* (Gobiesocidae) provides an intriguing system for testing these genomic trajectories. Unlike other clingfishes, *Gouania* species are highly specialized for an interstitial lifestyle in intertidal gravel beaches, where adults have extremely limited mobility (Briggs, 1955; Wagner et al., 2023). Connectivity is therefore thought to depend mainly on a short pelagic larval phase (∼12 days), probably with dispersal largely constrained to nearshore environments (Macpherson & Raventos, 2006). Together with the patchy distribution of suitable gravel beach habitats, this life history should promote fine-scale population differentiation (Wagner et al., 2019, 2021).

The genus comprises five species that diversified after the Messinian salinity crisis and whose distributions broadly align with major Mediterranean ecoregions (Figure 1a) (Wagner et al., 2019, 2021). One species, *G. willdenowi*, is restricted to the western Mediterranean, whereas the remaining four occur in the Adriatic and eastern Mediterranean. Importantly, these include two sympatric species pairs: *G. adriatica* and *G. pigra* in the Adriatic and *G. orientalis* and *G. hofrichteri* in the eastern basin. This combination of restricted distributions, replicated sympatry, ecological similarity and low adult dispersal makes *Gouania* particularly suitable for asking whether shared seascapes generate predictable patterns of connectivity – and whether such predictability extends to genomic landscapes of divergence.

**Figure 1.**
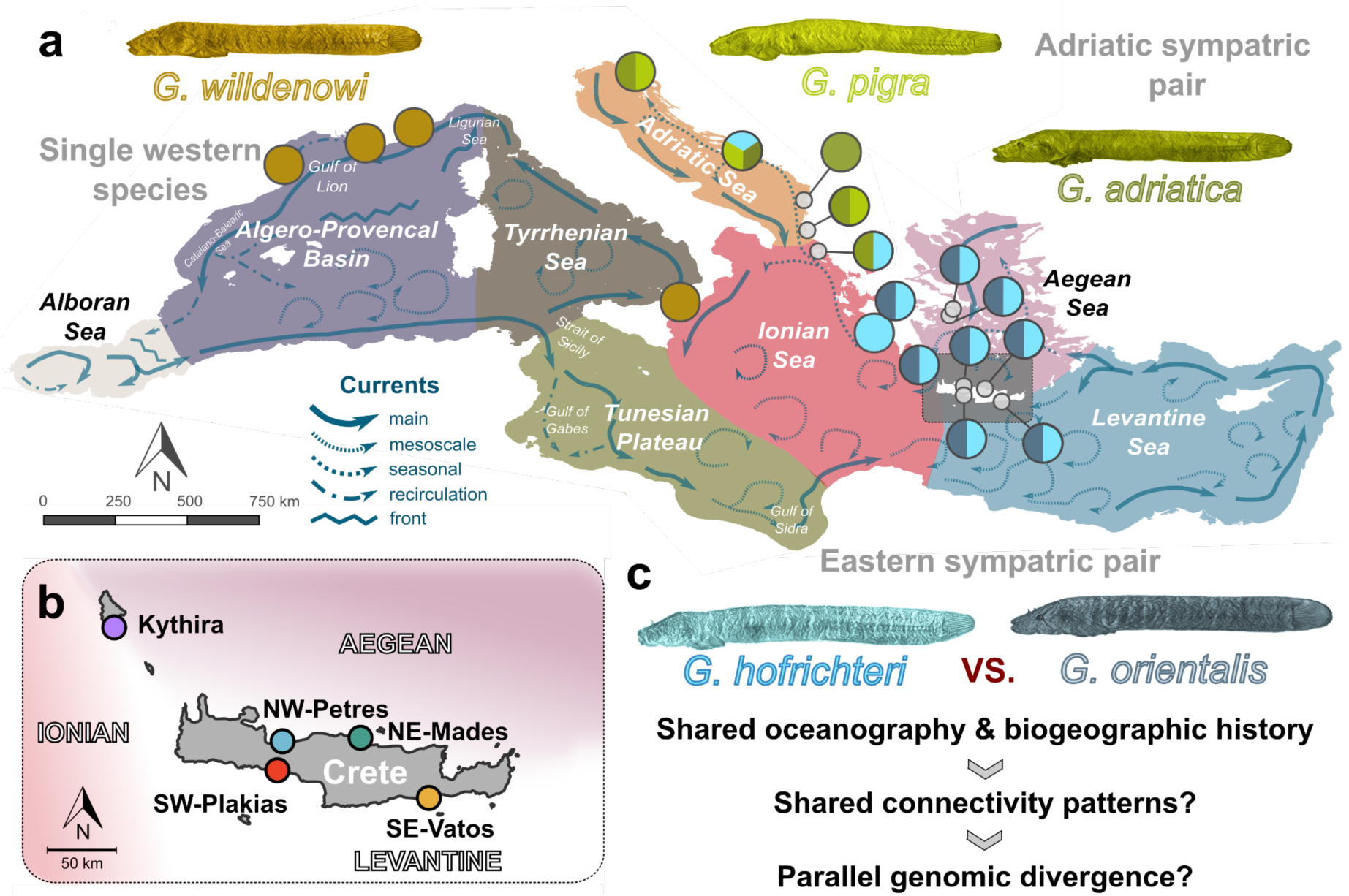
Biogeographic context and sampling design around the island of Crete in Mediterranean *Gouania*. **(a)** Distribution ranges and sampling sites of the five *Gouania* species, shown in different colors. Sympatric species pairs are indicated by half-filled circles. The background map is colored according to a Mediterranean biogeographic model comprising seven major ecoregions (sensu Spalding et al., 2007) and includes the major surface circulation patterns (El-Geziry & Bryden, 2010). Species illustrations are shown above their respective distribution ranges. **(b)** Sampling localities and **(c)** studying design around the island of Crete and Kythira. The study region is located at the intersection of three Mediterranean ecoregions (Ionian, Aegean and Levantine Seas) and therefore provides an ideal natural setting to investigate how shared oceanographic conditions influence connectivity and genomic divergence. Comparative whole-genome analyses focused on the two sympatric eastern Mediterranean species, *G. orientalis* and *G. hofrichteri*.

We tested this on a sympatric species pair *Gouania orientalis* and *G. hofrichteri* around Crete and the nearby island of Kythira. Crete is geographically isolated from the mainland, yet embedded in a highly structured circulation regime shaped by major jets and gyre systems and positioned at the intersection of the Ionian, Aegean and Levantine ecoregions (El-Geziry & Bryden, 2010; Spalding et al., 2007; Figure 1b, 2a). This geography creates replicated population contrasts for the sympatric species *G. orientalis* and *G. hofrichteri* around Crete and the nearby island of Kythira. In this study, we use whole-genome population data from both species to test two predictions: first, that passive larval transport should generate concordant patterns of genomic connectivity; and second, that shared environmental or ecological contrasts may produce a detectable excess of parallel genomic divergence. We then place these genomic results into a broader phylogeographic context using mitochondrial COI variation across all five *Gouania* species sampled throughout the Mediterranean basin.

## METHODS

### Whole genome re-sequencing and variant calling of sympatric *Gouania orientalis* and *G. hofrichteri* around Crete

For the two eastern Mediterranean sympatric species, *Gouania orientalis* and *G. hofrichteri*, we re-sequenced whole genomes from four populations around the island of Crete and from one population of the island of Kythira, altogether leading to 46 genomes of *G. orientalis* and 39 genomes of *G. hofrichteri* (Table S1). By focusing on co-occurring species within the same oceanographic context, we aimed to disentangle shared, environmentally driven patterns of connectivity from species-specific demographic and evolutionary responses (Figure 1b).

We sequenced most individuals to a target coverage of 5X but also two individuals per population to 15X coverage (Table S1). We aligned all raw reads against the *Gouania adriatica* reference genome (fGouWil2.1; (Rhie et al., 2021)) using BWA v0.7.17 and extracted variants with bcftools v1.14 (Danecek et al., 2021). We filtered sites with mapping quality of zero for more than 10% of reads or root mean square mapping quality of less than 50. We further masked sites with significant (p < 0.001) strand difference and with an extremely low (<2.5 percentile) or high (*>*97.5 percentile) depth. We filtered heterozygous sites exhibiting significant allelic depth imbalance between reference and alternative alleles using a binomial test (PHRED score >20), affecting between 0.02% and 0.15% of heterozygous sites per individual. Further, we filtered out sites with excess heterozygosity (InbreedingCoeff < 0.2) and sites with more than 20% missing genotypes. Our final variant call sets for *G. orientalis* and *G. hofrichteri* included 5,822,506 and 3,173,859 biallelic SNPs and indels, respectively.

### Lagrangian simulations around Crete

To test for correlations between oceanography and population structure for the population samples of *G. orientalis* and *G. hofrichteri* around the islands of Crete and Kythira as a null model for purely passive dispersal, we conducted Lagrangian simulations. We used the parcels package in python v3.1.1 (Delandmeter & Van Sebille, 2019; Lange & Van Sebille, 2017) based on hourly 2D current velocity data with a spatial resolution of 4.2 km from 01.11.2022 to 05.02.2025 downloaded from the Copernicus Marine Service, Mediterranean Sea Physics Analysis and Forecast (Clementi et al., 2023). Due to the coarse spatial resolution, the starting positions were not located within the current raster data. Therefore, we adapted the start position for each population for the larval dispersal modelling to the nearest raster cell centroid and used this as the new start position with longitude and latitude values (separated by comma) as follows: i) Kythira: 22.91666603088379, 36.14583206176758; ii) Petres 24.375, 35.39583206176758; iii) Mades 25.04166603088379, 35.4375; iv) Plakias 24.375, 35.14583206176758; and v) Vatos 25.58333396911621, 34.97916793823242. Throughout the whole period, we released one propagule per site each day using the fourth-order Runge-Kutta integration, following a rough estimation for the pelagic larval duration (PLD) of *G. willdenowi* of 13 days (Macpherson & Raventos, 2006) and recorded the location of each larva every 6 hours. Due to a lack of knowledge about the breeding season for the investigated species, we assessed seasonal drift differences between spring (March-June), summer (June-September), autumn (September-November) and winter (November-March) and we conducted a second simulation based on a PLD of 26 days.

### Fine-scale genomic population structure and connectivity analyses around Crete

We then tested for gene flow and connectivity among populations calculating *D*-statistics in Dsuite v0.4 r42 (Malinsky et al., 2021) using the Dtrios function. For the latter, we included an individual of the other species (*G. orientalis* or *G. hofrichteri*) as an outgroup in the respective individual VCF file and adjusted the *p*-values for multiple testing by applying a Benjamini-Hochberg correction (Benjamini & Hochberg, 1995). In order to assess population genomic metrics, we calculated genome-wide mean estimates of nucleotide diversity (*pi*), Fixation index (*F_ST_*, (Hudson et al., 1992) and mean net-between group divergence (*d_A_*) using sliding windows of 60,000 bp with a step size of 30,000 bp. Windows containing fewer than 100 SNPs were excluded. Calculations were performed using the popgenWindows.py script from https://github.com/simonhmartin/genomics_general/tree/master and a standard calculation for *d_A_* (following calculations and codes in (Wagner et al., 2026). We also calculated the average genome wide individual inbreeding coefficient (*F_IS_*) using vcftools 0.1.16 (Danecek et al., 2011). To quantify the similarity of population differentiation patterns between species, we compared mean pairwise *F_ST_* estimates for all population pairs using Spearman’s rank correlation. Correlations were calculated from the corresponding pairwise *F_ST_* values derived independently for *G. orientalis* and *G. hofrichteri*.

For inferring phylogenetic clusters, we created a distance matrix using PLINK v1.90b6.21 (Purcell et al., 2007). Principal component analyses (PCA) were performed using PLINK2 v2.0 on genome-wide SNP datasets converted to binary PLINK format. PCA was conducted using the `--pca` function following filtering for minor allele frequency (MAF) thresholds of 0.01, 0.05, 0.10, 0.20 and 0.40. Eigenvectors and eigenvalues were extracted from the resulting covariance matrix and used for downstream visualization for MAF of 0.1. This led to 1,071,073 and 1,054,075 biallelic SNPs, respectively, for *G. orientalis* and *G. hofrichteri*. We investigated population genetic structure using PCA in PLINK using a MAF of 0.15 and model-based ancestry estimation with ADMIXTURE v1.3.0 (Alexander et al., 2009).

### Local (parallel) genomic signals of population-specific divergence

We annotated variants from the VCF with SnpEff (Cingolani et al., 2012), using the fGouWil2.1.99 gene annotation. To improve annotation of unresolved Ensembl gene identifiers (ENSGWIG) in the fGouWil2.1.99 genome annotation, a custom annotation table was generated by combining Ensembl and NCBI gene annotations. Remaining unresolved protein-coding genes were annotated by retrieving canonical protein sequences from Ensembl and searching them against the UniProt/Swiss-Prot database using DIAMOND BLASTP (Buchfink et al., 2021). For each query, the best-scoring Swiss-Prot hit was retained and gene symbols were assigned for alignments with ≥50% query coverage and an E-value < 1 × 10⁻⁵. Existing annotations were retained and Swiss-Prot-derived annotations were only used for previously unresolved genes. This procedure provided annotations for an additional 6,554 protein-coding genes and improved the functional annotation used in downstream analyses.

First, we specifically focused on reciprocal patterns, where alleles were close to fixation in Kythira but rare or absent in mainland Crete populations, or vice versa (AF ≥ 0.95 or AF ≤ 0.05). To assess whether parallelism was also detectable at the gene level among these nearly fixed variants, we performed a permutation test in which gene sets were randomly sampled from the background set of 23,789 annotated genes while preserving the observed number of genes containing nearly fixed variants in G. orientalis and G. hofrichteri (45 and 8 genes, respectively). Significance was calculated as the proportion of permutations yielding gene overlap equal to or greater than that observed between species.

We then used two related but complementary approaches to identify genomic outlier windows. The Population Branch Statistic (PBS) (Yi et al., 2010) captures lineage-specific differentiation in the focal population while controlling for shared ancestry among populations, whereas *dA* identifies genomic windows with high relative divergence. To identify robust PBS outliers reflecting lineage-specific differentiation in Kythira, we calculated PBS for Kythira (PBS_Kythira_) across multiple trio configurations involving Kythira and pairs of Cretan populations. For each trio-specific PBS distribution, the 95th percentile was calculated across all genomic windows. A window was classified as a PBS outlier only if its PBS value for Kythira exceeded the 95th percentile threshold in all trios, ensuring that inferred signals of branch-specific differentiation were consistent across alternative reference population combinations. Second, for each pairwise comparison between Kythira and a Cretan population, *d_A_* outliers were defined as genomic windows falling within the 95th percentile of the *d_A_* distribution. To account for background genetic structure within Crete, *d_A_* windows that were also identified as outliers in any Crete-Crete comparison were excluded. Like the PBS approach, this analysis was intended to account for biases associated with population structure within the Cretan lineage, focusing on genomic regions that consistently distinguish Kythira from the broader Cretan lineage. By intersecting PBS and *d_A_* outliers, we defined a set of high-confidence genomic windows associated with population-specific divergence in Kythira.

We then identified parallel genomic regions as those windows classified as outliers in both species under the same criteria. To assess whether overlap among candidate windows exceeded random expectations, we performed a hypergeometric test using the total number of genomic windows analysed as the background set (29,039 windows), while preserving the observed number of candidate windows in each species. Expected overlap and enrichment were calculated relative to the null expectation of random overlap. To further evaluate whether these candidate windows showed signatures consistent with selection, we calculated Tajima’s D (Tajima, 1989) within Kythira. We then tested whether candidate windows were significantly enriched for Tajima’s D values more negative than the genomic background, consistent with recent selective sweeps. To facilitate direct comparisons between populations and control for non-population specific evolutionary processes, we calculated ΔTajima’s D as the difference between Tajima’s D in Kythira and the mean Tajima’s D across Crete populations for each genomic window.

We then extracted candidate genes from these outlier windows from the *Gouania adriatica* annotation (fGouWil2.1.115). For each outlier window, we extracted genes overlapping a ±30 kb interval around the window midpoint on the same scaffold.

Finally, we conducted three functional enrichment analyses performed with g:Profiler (Reimand et al., 2007) using gene sets unique to (i) *G. orientalis*, (ii) *G. hofrichteri* and (iii) the overlap of windows between the two species (i.e., parallel outlier windows). This approach allowed us to investigate both independent and shared (parallel) genomic pathways underlying local adaptation. All GO enrichment analyses were performed using zebrafish GO annotations with a custom gene background (all *Gouania adriatica* reference genes) applying an FDR threshold of 0.05 and a Benjamini-Hochberg correction (Benjamini & Hochberg, 1995).

### Mediterranean basin-wide phylogeographic reconstructions based on COI data

For assessing phylogeographic patterns across the genus, we analysed a total of 668 partial sequences of the mitochondrial cytochrome-c-oxidase subunit I gene (COI) from 23 populations, generated in previous studies (Wagner et al., 2019, 2021, 2026) and deposited on GenBank (accession numbers: MK873443–MK873539, MT299844–MT299870 and OL839338–OL944591 summarized in Table S2). We aligned and trimmed the individual COI sequences in MEGA 7.0 (Kumar et al., 2016) and applied the integrated MUSCLE (Edgar, 2004) algorithm. We visualized the geographic structure of intraspecific mitochondrial genetic diversity for each of the five *Gouania* species by means of a parsimony network using TCS (Clement et al., 2002) in PopART v.1.7 (Leigh & Bryant, 2015).

### Demographic analysis based on whole genome data

For inferring the demographic histories of individual *Gouania* species, we applied the Multiple Sequentially Markovian Coalescent (MSMC2) model (Schiffels & Wang, 2020) to a deep-sequenced (∼30-fold coverage) specimen of each *Gouania* species (*G. hofrichteri* and *G. orientalis* from Attica, Greece; *G. adriatica* as well as *G. pigra* from Pula, Croatia and *G. willdenowi* from Toulon, France). The individual raw sequence files were generated in a previous study (Wagner et al., 2026) and accessed from the European Nucleotide Archive using the numbers: ERS9744577 (*G. willdenowi*; Toulon, France), ERS9744566 (*G. pigra*; Pula, Croatia), ERS9744553 (*G. orientalis*; Saronida, Greece) and ERS9744540 (*G. hofrichteri;* Chamolia, Greece). As we were lacking this data for *G. adriatica*, we used short-read data obtained for the creation of the reference genome (Rhie et al., 2021), by downloading a fraction of the original raw data (“fGouWil2_S1_L007_R[1&2]_001.fastq.gz”; Pula, Croatia) from GenomeArk (https://genomeark.github.io), which yielded about the same coverage as individuals from the other species. We visualized the results assuming a mutation rate of 3.5e-08 (Malinsky et al., 2018) and an estimated generation time of 2 years (own aquarium observation). For each MSMC2 run we performed 20 bootstrap replicates following the method and plotting scripts from (Lescroart et al., 2023).

### Open Research and Data Availability

All bioinformatic analyses were implemented using workflow-based pipelines developed in Snakemake v7.25.0 (Mölder et al., 2021), and downstream analyses, visualization, and statistical analyses were conducted in Jupyter notebooks. The complete analysis workflows and scripts used in this study are publicly available at GitHub (https://github.com/maxwagn/GouaniaPhylogeographyCrete) and, for the particle-drifting simulations, at https://github.com/arianedr/Particle_Simulation/blob/main/Particle_Drifting.ipynb. Archived versions of all local analysis scripts, notebooks, intermediate files, and supplementary datasets are available through Figshare (https://doi.org/10.6084/m9.figshare.32686272).

Raw whole-genome sequencing data generated for this study have been deposited in the European Nucleotide Archive (ENA) under accession ERR7917043–ERR7917126 (Table S1). Mitochondrial COI sequences analysed in this study are publicly available through GenBank under accession numbers MK873443–MK873539, MT299844–MT299870, and OL839338–OL944591 (Table S2).

## RESULTS

### Lagrangian simulations predict structured passive dispersal around Crete

To generate explicit predictions for passive larval transport around the island of Crete, we performed Lagrangian passive particle drift simulations (Figure 2). Assuming a drifting duration of ∼13 days, corresponding to previous estimates of pelagic larval duration in *Gouania* (Macpherson & Raventos, 2006), simulations revealed pronounced spatial structure across all seasons (Figure 2b–e).

**Figure 2.**
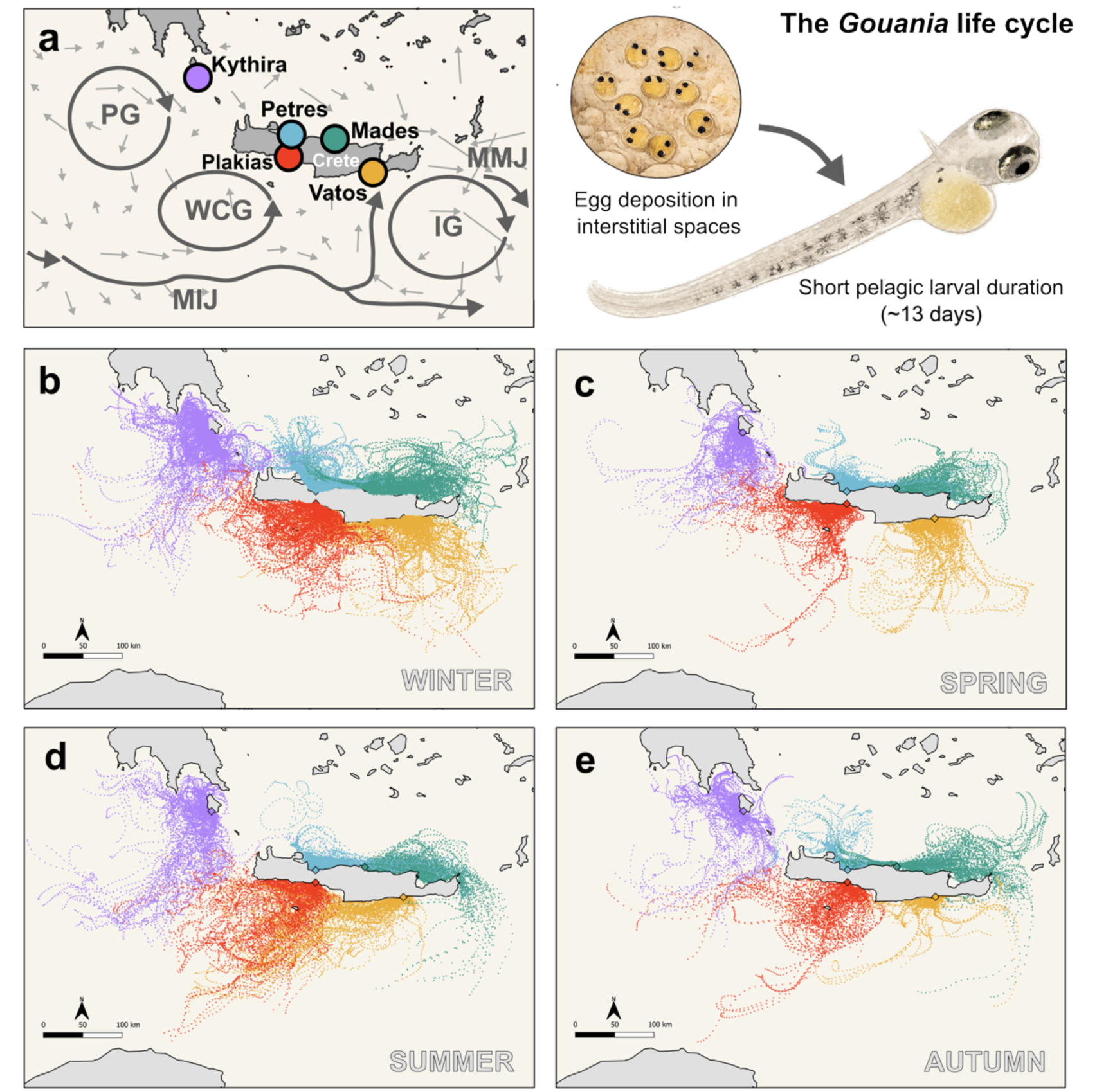
Passive larval drift simulation as a null model for dispersal abilities in *Gouania*. **(a)** The five investigated populations are embedded in a complex seascape with three main gyres and jets: the Pelops Gyre (PG), the Western Cretan Gyre (WCG), the Ierapetra Gyre (IG), the Mid-Ionian Jet (MIJ) and the Mid-Mediterranean Jet (MMJ). Arrows represent mean currents in 0.5° x 0.5° grids This map was adapted from published drift data (Poulin et al., 2013). **(b–e)** Lagrangian simulations of passive larval drift around Crete and Kythira across four seasons: **(b)** winter (November–March), **(c)** spring (March–June), **(d)** summer (June–September) and **(e)** autumn (September–November). Particles were released daily from the four sampled populations (Kythira, NW-Petres, NE-Mades and southern Crete) and tracked for 13 days. Each point represents particle positions at 6-hour intervals.

Kythira was largely isolated from Crete and northern and southern Cretan populations were strongly separated. Dispersal trajectories showed substantial overlap between the two northern populations, Mades and Petres, across all seasons. In contrast, connectivity between the southern populations, Vatos and Plakias, was weaker, particularly during spring (Figure 2c). Seasonal simulations further suggested limited and asymmetric exchange around Crete. During summer (Figure 2d), a small proportion of particles released from northern Crete, particularly Mades, reached the southern coastline, whereas in autumn (Figure 2e), limited northward dispersal from southern Crete, especially Vatos, was observed.

Overall, the simulations predict a spatially structured and seasonally variable dispersal regime, with strongest connectivity among northern populations, limited exchange between northern and southern Crete and weak connectivity between Crete and Kythira. The trajectories also suggest a predominantly clockwise pattern of potential passive dispersal around Crete. Using a longer drifting duration of 26 days yielded largely consistent results (Figure S1), suggesting that these predictions are not particularly sensitive to the assumed pelagic larval duration. In the following section, we evaluate this null model of dispersal driven solely by physical oceanographic processes against genome-wide population structure.

### Whole-genome data reveal concordant connectivity patterns in sympatric *Gouania*

To test whether the passive-dispersal predictions were reflected in realised genomic connectivity, we analysed whole-genome population structure in the sympatric species *Gouania orientalis* and *G. hofrichteri* around Crete and Kythira. Across both species, we found a pronounced division between northern and southern Cretan populations, with individuals from Kythira clustering more closely with northern Crete than with southern Crete. This pattern was consistently supported by PCA, pairwise *F*_ST_, admixture analyses and tests of excess allele sharing (Figure 3; Figures S2–S3; Table 1).

**Figure 3.**
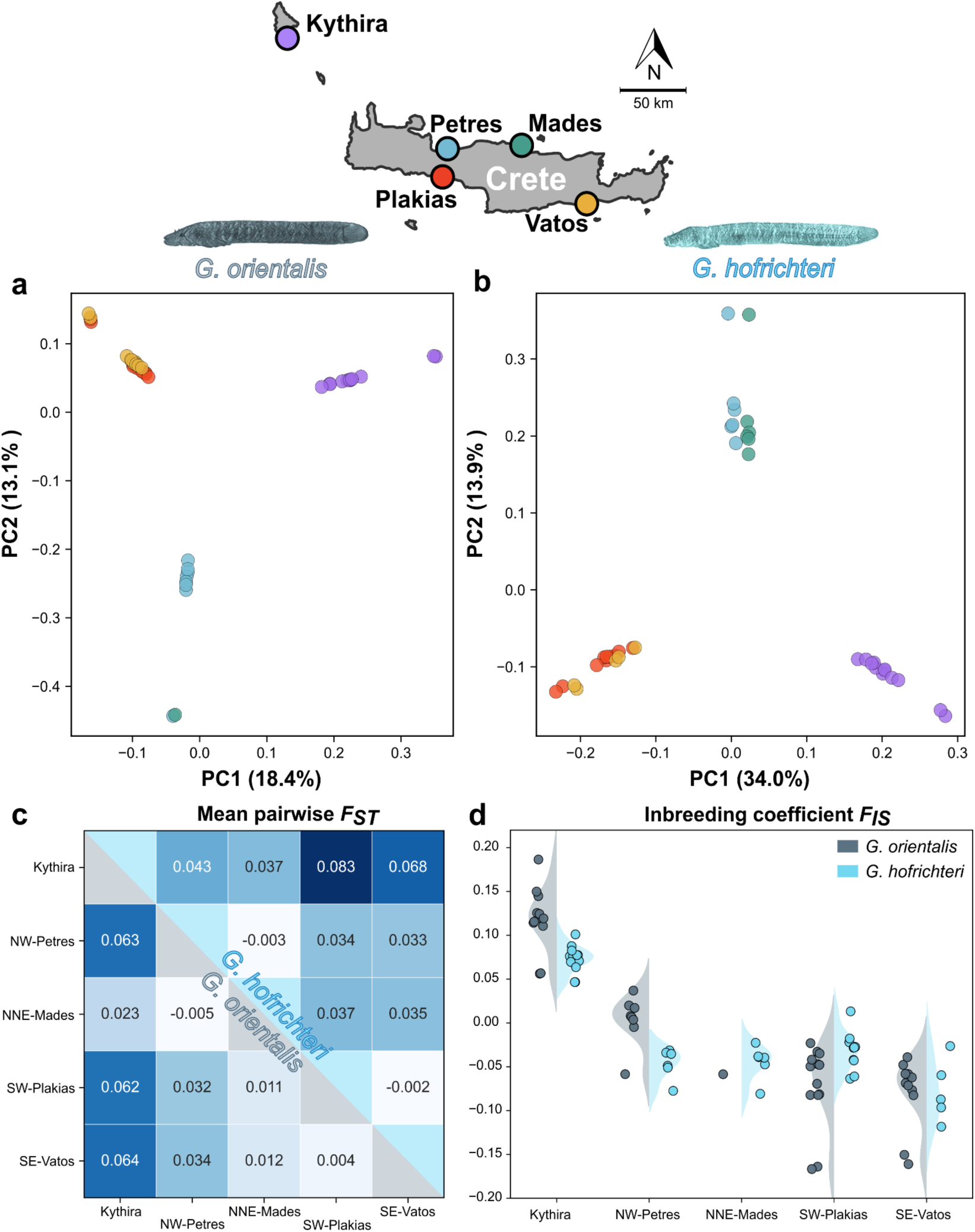
Parallel population genomic structure across sympatric *Gouania* species around the island of Crete. Principal components analysis of biallelic SNPs (MAF=0.1) for *G. orientalis* **(a)** and *G. hofrichteri* **(b)**. Colored dots correspond to sampling locations shown on the map above. Roman numerals (I–III) indicate regions of increased network complexity. **(c)** Pairwise genome-wide mean *F_ST_* between populations for *G. orientalis* (lower diagonal) and *G. hofrichteri* (upper diagonal). **(d)** Distribution of genome-wide inbreeding coefficients (*F_IS_*) across populations for both species. Elevated *F_IS_* values in Kythira are consistent across species, suggesting increased inbreeding and/or stronger population subdivision relative to the Cretan populations.

**Table 1.**
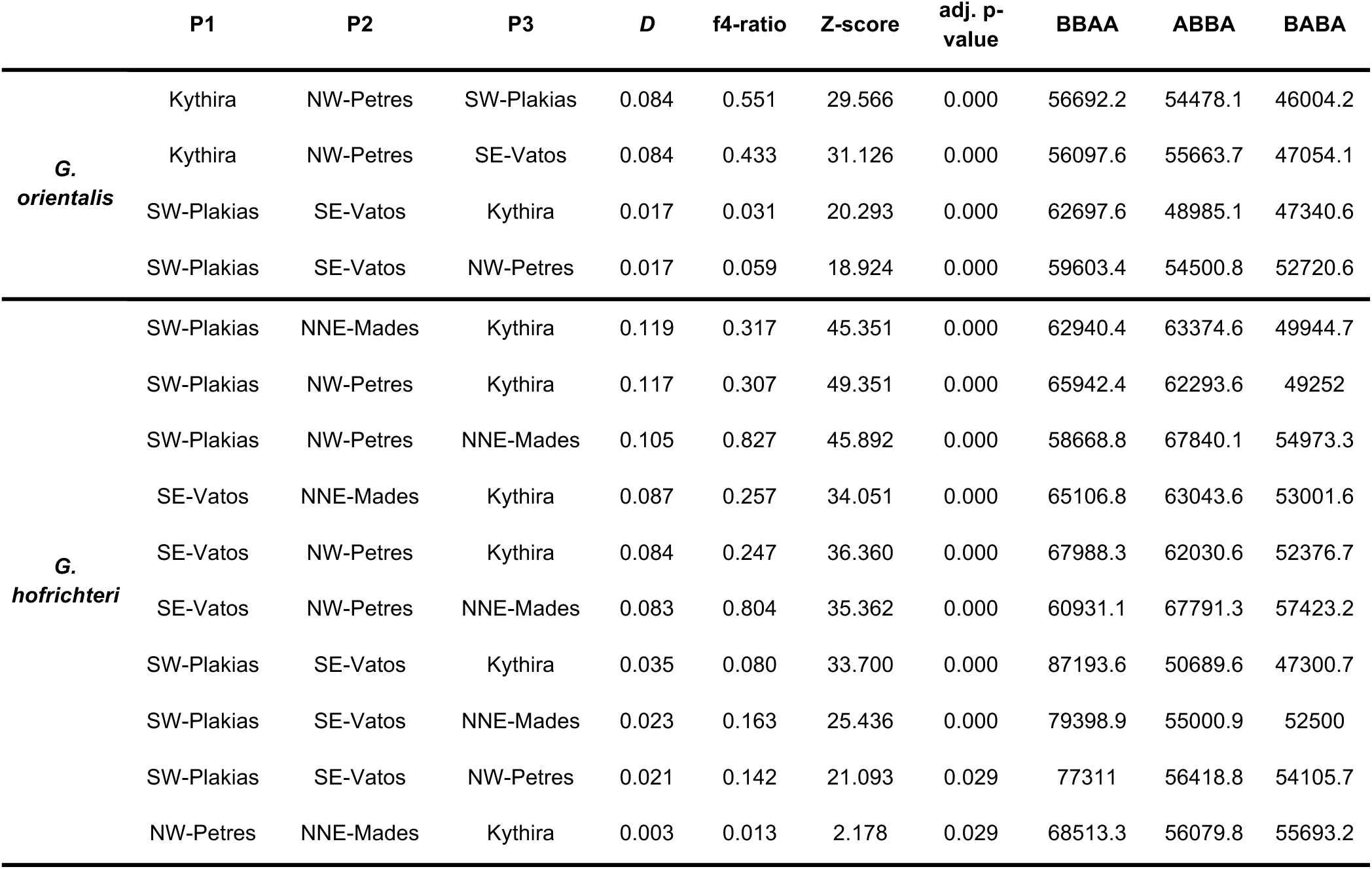
Population connectivity among *Gouania orientalis* and *G. hofrichteri* populations around the island of Crete and Kythira. D- and f4-statistics were calculated with Dsuite (P1-P3 sorted by phylogeny). *p*-values were adjusted for multiple testing applying a Benjamini-Hochberg correction and significant trios are presented in bold. For the *G. orientalis* dataset, we excluded the single population sample from NE-Mades as it led to biases in the statistics.

Principal component analyses revealed broadly congruent population structure in both species (Figure 3a,b). In both cases, PC1 separated Kythira from the Cretan populations, whereas PC2 captured a north–south subdivision within Crete. In *G. orientalis*, the single individual from NNE-Mades grouped closely with NW-Petres. In *G. hofrichteri*, the north–south subdivision was also evident, but population clustering within Crete was more pronounced. Together, these results indicate shared geographic structure across species, characterized by strong differentiation of Kythira and a secondary north–south break within Crete.

Admixture analyses supported these patterns. In *G. orientalis*, Kythira was strongly differentiated from the Cretan populations (Figure S2a). In contrast, northern Cretan populations of *G. hofrichteri* showed admixture components also present in both Kythira and southern Crete (Figure S2b), consistent with connectivity mediated stepping-stone-wise through northern Crete. When analyses were restricted to Cretan populations only, both species showed a pronounced north–south structure (Figure S3).

Patterns of population differentiation were highly concordant between species. Pairwise *F_ST_* estimates were strongly correlated between *G. orientalis* and *G. hofrichteri* across all population comparisons (Spearman’s ρ = 0.733, *P =* 0.0158), indicating broadly similar patterns of genetic structure (Figure 3c). In both species, differentiation was low among populations from the same side of Crete and generally higher between northern and southern populations. Comparisons involving Kythira showed the highest levels of differentiation. In *G. hofrichteri*, Kythira was less differentiated from northern than from southern Cretan populations, whereas differentiation between Kythira and Crete was more uniform in *G. orientalis*. Comparisons involving the NNE-Mades population of *G. orientalis* yielded consistently low *F_ST_* values, but this pattern should be interpreted cautiously because only a single individual was sampled from this locality.

Tests of excess allele sharing largely mirrored the connectivity patterns predicted by passive larval drift simulations. In both species, D-statistics revealed significant excess allele sharing among northern and southern Cretan populations (Table 1), despite the inferred sister relationship between Kythira and northern Crete in the population trees. In particular, excess allele sharing between SE-Vatos and northern Cretan populations relative to SW-Plakias suggests clockwise connectivity around the eastern flank of Crete, consistent with the simulated dispersal trajectories.

Both species showed a similar relative pattern, with Kythira exhibiting significantly lower nucleotide diversity and higher inbreeding than other populations (Figure 3d; Figure S4). Individual inbreeding coefficients (*F_IS_*) differed strongly among populations (two-way ANOVA, population effect: *F_4,75_* = 98.62, *P =* 4.37 × 10⁻²⁹), whereas the overall effect of species was not significant (*F_1,75_*= 2.23, *P =* 0.140). However, a significant species × population interaction (*F_4,75_* = 6.72, *P =* 1.12 × 10⁻⁴) indicated that population-specific patterns of inbreeding differed between *G. orientalis* and *G. hofrichteri*. Genome-wide diversity metrics revealed both species-specific differences and shared population-level trends. Nucleotide diversity (π) was consistently higher in *G. orientalis* than in *G. hofrichteri* across populations (Figure S4). Despite this difference, both species exhibited remarkably similar spatial patterns of diversity. Kythira showed the lowest nucleotide diversity in both species, whereas the Cretan populations displayed comparatively higher and more homogeneous levels of diversity. The single *G. orientalis* individual from NE-Mades showed considerably higher nucleotide diversity than other populations, which likely contributed to the lower *F*_ST_ values observed in comparisons involving this locality.

### Shared population contrasts reveal parallel and species-specific genomic divergence

The concordant population structure observed in *G. orientalis* and *G. hofrichteri* around Crete provides an opportunity to test whether shared population contrasts are associated with parallel patterns of genomic differentiation across the genome. In both species, Kythira was strongly differentiated from Crete, showed reduced nucleotide diversity and had elevated inbreeding coefficients, making it a comparable focal population across two independently evolving lineages. We therefore used Kythira-centred comparisons to ask whether divergence between Kythira and Crete involved the same genomic regions, genes, or pathways in both species, as expected if some differentiation outliers reflect local adaptation driven by shared selective pressures.

To identify genomic regions associated with Kythira-specific divergence (Figure 4; Figure S5), we quantified PBS and *d_A_* outliers across 29,039 genomic windows. In *G. orientalis*, 966 windows exceeded the genome-wide 95th percentile for PBS across all three Kythira-centred trios. In *G. hofrichteri*, 347 windows met this criterion across six Kythira-centred trios. This difference partly reflects the exclusion of the single NE-Mades individual from the *G. orientalis* analysis, which reduced the number of informative Kythira-centred trios relative to *G. hofrichteri*. Using *d_A_*, we detected 1,612 Kythira-specific outlier windows in *G. orientalis* and 2,276 in *G. hofrichteri*.

**Figure 4.**
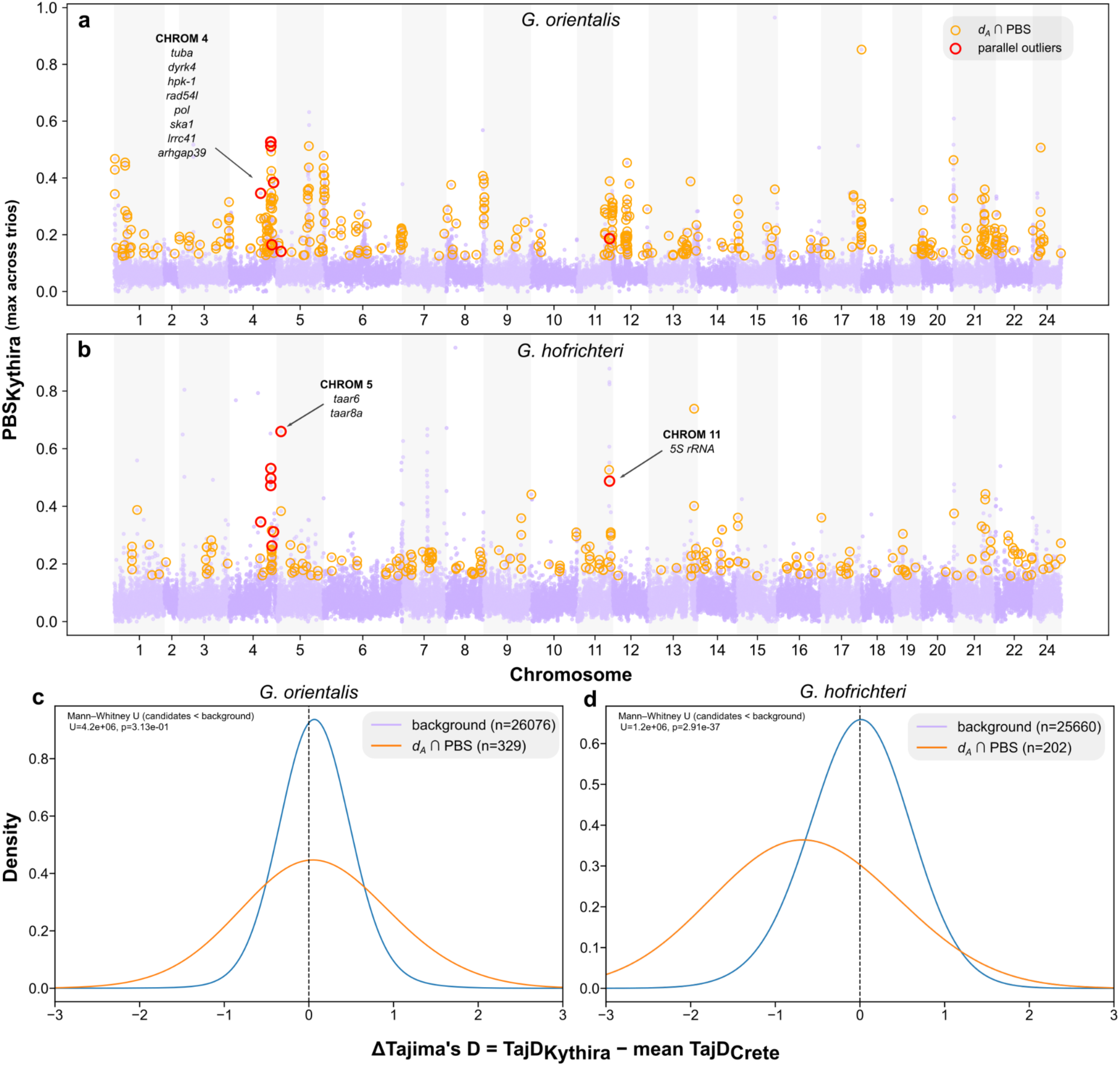
Parallel and species-specific genomic signals between sympatric *Gouania orientalis* and *G. hofrichteri* Kythira and Crete populations. Maximum PBS values for the Kythira population, summarized across all trio comparisons, are shown for (a) *G. orientalis* and (b) *G. hofrichteri*. Orange-outlined points indicate genomic windows identified as outliers by both *d_A_* and PBS, while red circles highlight parallel outlier windows shared between the two species. Arrows indicate genes found within the surrounding area of outlier windows (a full list is shown in Table S6). Furthermore, density distributions of ΔTajima’s D (Tajima’s D_Kythira_ − mean Tajima’s D_Crete_) compare candidate windows against the genomic background, revealing no significant shift towards more negative values in (c) *G. orientalis*, but a significant enrichment of more negative values in (d) *G. hofrichteri*.

We then defined final candidate windows as those identified by both PBS and *d_A_*. This yielded 397 candidate windows in *G. orientalis* (Figure 4a; Figure S5a) and 220 candidate windows in *G. hofrichteri* (Figure 4b; Figure S5b), whereas only 8 were shared between the two species, (i.e., parallel outlier windows). This overlap was significantly higher than expected under random overlap: 3.01 windows were expected, corresponding to a 2.66-fold enrichment (hypergeometric test, *P* = 0.011). These shared windows therefore represent a conservative set of parallel genomic regions associated with the Kythira–Crete contrast in both species.

At the gene level, these 397 candidate windows in *G. orientalis* and 220 candidate windows in *G. hofrichteri* corresponded to 895 and 656 unique genes, respectively (Table S3). Intersecting these sets identified 50 shared genes, which is significantly more than expected by chance (permutation test, *P* = 4.0 × 10⁻⁵), providing further evidence for parallel genomic divergence associated with the Kythira–Crete contrast on a gene-level.

We further examined whether parallelism was detectable at the level of nearly fixed variants. We identified 54 and 12 nearly fixed variants (SNPs and indels) associated with 52 and 14 genes in *G. orientalis* and *G. hofrichteri*, respectively (Table S3). Although no identical nearly fixed variants were shared between species, both contained Kythira-specific nearly fixed variants associated with the gene *tuba* (Tables S4, S5). This overlap was highly unlikely to arise by chance given the observed numbers of candidate genes (permutation test, *P* = 0.0334). These results therefore suggest parallelism at the gene level involving *tuba*, despite the absence of shared underlying variants.

We next assessed whether candidate regions showed signatures consistent with recent or localised selection by examining Tajima’s D. In contrast to *G. orientalis* (Figure 4c), candidate windows in *G. hofrichteri* were significantly shifted towards more negative Tajima’s D values relative to the genomic background (Mann–Whitney U; *P* = 2.91 × 10^-37^), indicating a Kythera-specific excess of low frequency variants in these regions and consistent with an enrichment for population-specific selective sweeps in Kythira (Figure 4d). At the genome-wide scale, however, *G. orientalis* showed generally more negative Tajima’s D values across all populations, which may influence the detectability of localised selection (Figures S6, S7).

Finally, we tested whether candidate genes were enriched for specific functional categories. GO enrichment analyses of genes located in parallel windows and overlapping outlier windows revealed no significant enrichment. Likewise, species-specific candidate gene sets yielded predominantly broad and non-specific functional categories (Table S7, S8). Outlier-associated genes in both species were enriched for general molecular functions and cellular processes, including ion binding, protein binding, catalytic activity and metabolic pathways. In *G. hofrichteri*, enriched terms were primarily associated with phosphorylation and phosphorus-containing compound metabolism, whereas *G. orientalis* showed additional enrichment for protein modification, cell motility and cadherin signalling. Overall, enrichment patterns were diffuse and did not identify a single dominant biological pathway, suggesting that adaptive divergence may involve numerous genes with diverse cellular functions rather than a small number of functionally coherent pathways.

### Genus-wide mtDNA variation reveals broad basin-level structure but species-specific phylogeographic patterns

Across the genus, mitochondrial COI variation revealed pronounced differences in phylogeographic structure among *Gouania* species (Figure 5). The western Mediterranean species, *G. willdenowi* and the two Adriatic species showed comparatively compact haplotype networks. *G. willdenowi* exhibited a shallow, star-like network dominated by a common haplotype shared among French populations, with the Messina population represented by a private but closely related haplotype (Figure 5a).

**Figure 5.**
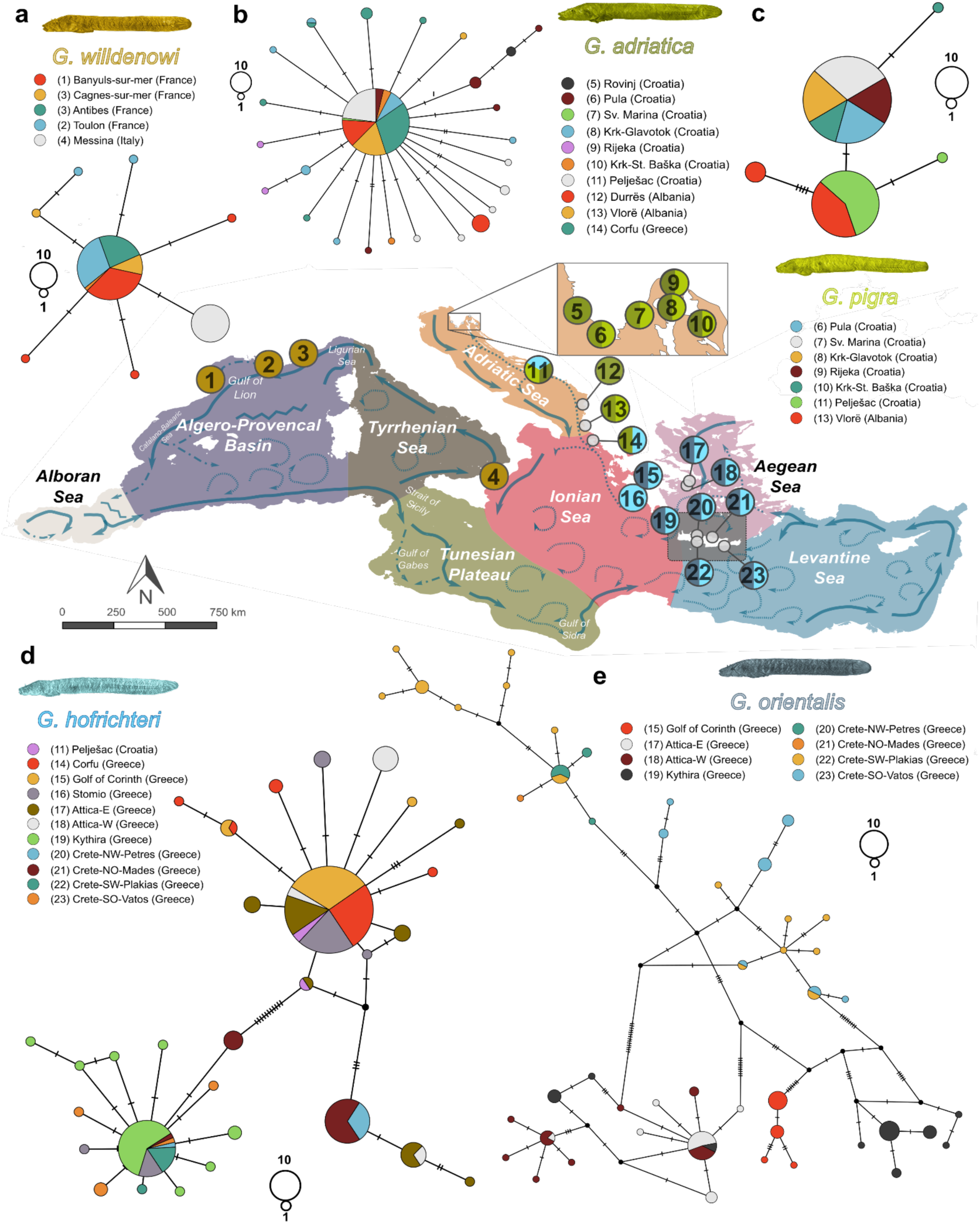
Bio-and phylogeographic patterns based on COI data of all five closely related beach fishes of the genus *Gouania*. (**a-e)** TCS haplotype networks are based on partial COI sequences for each of the five *Gouania* species. For each network two circles representing the size of ten and one sample are given. The numbers before each location name in brackets refers to the same numbers in grey circles corresponding to sampling sites shown in the centered map. Note that because of their vicinity, populations from Canges-sur-mer and Antibes show the same number in the map. For a detailed list of samples per population see Table S2.

Similarly, *G. adriatica* showed a star-like network dominated by a common central haplotype (Figure 5b), whereas *G. pigra* showed two main haplotype groups corresponding broadly to northern and southern Adriatic populations (Figure 5c). These comparatively shallow networks are consistent with recent demographic expansions in western and Adriatic populations, possibly linked to Quaternary sea-level fluctuations and postglacial changes in shallow coastal habitat.

By contrast, the two eastern Mediterranean species, *G. hofrichteri* and *G. orientalis*, showed substantially stronger mitochondrial structure and higher haplotype diversity. However, this structure was not parallel at the level of individual populations. In *G. hofrichteri*, haplotypes from Crete and Kythira were largely associated with the same divergent mitochondrial cluster, separated from many mainland Greek and more northern populations (Figure 5d). In *G. orientalis*, mitochondrial structure was more complex and Kythira did not cluster with Crete but instead appeared closer to haplotypes from mainland/Aegean populations (Figure 5e).

### Species-level demographic histories are heterogeneous across *Gouania*

To obtain a species-level temporal perspective on demographic patterns across *Gouania*, we reconstructed effective population size (*N*e) trajectories with MSMC2. These analyses, based on one high-coverage individual per species, revealed substantial heterogeneity in inferred demographic histories across the genus (Figure 6).

**Figure 6.**
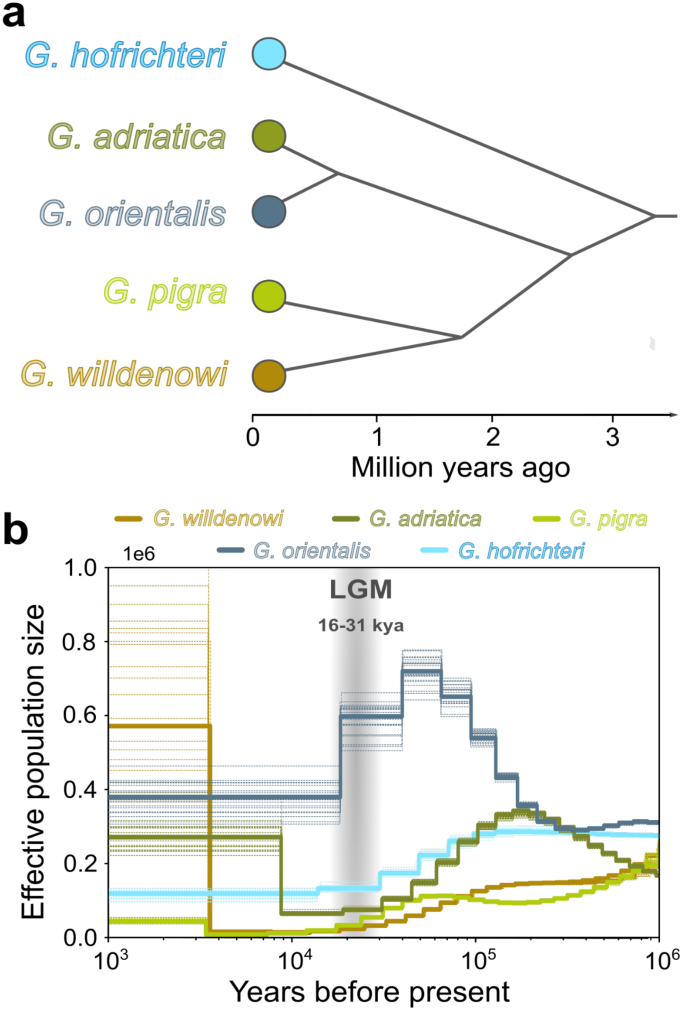
**Demographic histories inferred using MSMC2 for all five *Gouania* species**. **(a)** Post-Messinian radiation of the genus *Gouania*. Phylogenetic relationships among species are shown based on a simplified dated phylogeny inferred from nine nuclear and one mitochondrial marker (Wagner et al., 2019). **(b)** *G. orientalis* (Saronida, Attica, Greece), *G. hofrichteri* (Chamolia, Attica, Greece), *G. adriatica* (Pula, Croatia), *G. pigra* (Pula, Croatia) and *G. willdenowi* (Toulon, France). For each species, 20 bootstrap replicates are shown as dashed lines, with the main estimate represented by a solid line. Grey shading indicates the last glacial maximum (LGM) at around 20,000 years BP.

Overall, MSMC2 results indicate that species-level demographic histories are only partly concordant across *Gouania* and cannot be readily explained by either deep phylogenetic relationships (Figure 6a), present-day sympatry, or geographic co-occurrence. Instead, demographic trajectories appear to reflect a combination of deeper historical processes and species-specific responses to environmental change, contrasting with the stronger concordance observed for contemporary genomic connectivity around Crete.

The two eastern Mediterranean species showed distinct histories (Figure 6b). *Gouaniua orientalis* exhibited relatively high inferred *N*e through much of the late Pleistocene, followed by a decline towards more recent periods, whereas *G. hofrichteri* showed a lower trajectory over the same interval, with a decline between the late Pleistocene and the Last Glacial Maximum (LGM). Both species showed relatively stable *N*e since the LGM. The Adriatic species, *G. adriatica* and *G. pigra*, showed declines in inferred *N*e between the late Pleistocene and the LGM, followed by relative increases within the last 10 kyr, although their detailed trajectories and absolute *N*e values differed markedly. The western Mediterranean species, *G. willdenowi*, showed a trajectory broadly similar to its sister species *G. pigra*, with a sharper but more variable recent increase.

## DISCUSSION

A central assumption in comparative phylogeography is that closely related and ecologically similar species exposed to similar environmental conditions should exhibit comparable patterns of connectivity and divergence (Dawson, 2012). In this context, the sympatrically occurring species of the genus *Gouania* provide an ideal system for testing this hypothesis across both broad (Mediterranean-wide) and fine (within-region) spatial scales. Indeed, across multiple levels of biological organization, sympatric *Gouania* species exhibited a remarkable degree of concordance. Contemporary genomic connectivity around Crete closely matched oceanographic predictions in both species and shared population contrasts were associated with a significant excess of parallel genomic divergence. In contrast, mitochondrial phylogeographic patterns and species-level demographic histories were considerably more heterogeneous. Together, these results suggest that shared oceanographic and biogeographic contexts can generate predictable patterns of connectivity and aspects of genomic divergence, whereas deeper historical processes remain comparatively species-specific.

### Oceanography generates predictable connectivity patterns across sympatric species: insights from Crete

These patterns of predictability extend to smaller spatial scales, as demonstrated by our case study around the island of Crete. Both sympatric species, *Gouania orientalis* and *G. hofrichteri*, exhibit remarkably similar patterns of population structure that closely match the surrounding oceanographic regime. In particular, the pronounced differentiation between northern and southern Cretan populations, as well as the strong isolation of Kythira, are consistently recovered across both species and align closely with predictions from Lagrangian particle drift simulations.

This concordance between simulated passive dispersal and observed genomic connectivity indicates that oceanographic processes are likely a primary driver of gene flow in these species. Notably, this result challenges previous assumptions that *Gouania*, due to its short pelagic larval phase and interstitial lifestyle, might be largely restricted to nearshore dispersal (Macpherson & Raventos, 2006; Wagner et al., 2019, 2021). Instead, our results demonstrate that even limited pelagic durations (ca. 10–17 days) can be sufficient for oceanographic transport to structure population connectivity at regional scales. More generally, this is consistent with theoretical and empirical studies showing that even low levels of larval exchange can influence patterns of genetic structure in marine populations (Palumbi, 2003). At the same time, both species show elevated inbreeding coefficients in Kythira, consistent with strong local larval retention and suggesting that populations may persist as highly localized units embedded within weakly connected dispersal networks. Such nearshore retention is common in cryptobenthic fishes and is often linked to strong habitat specialization and settlement into spatially rare, patchily distributed microhabitats such as Mediterranean gravel beaches (Brandl et al., 2018; Wagner et al., 2019).

Despite these broadly parallel patterns, subtle but consistent differences between species indicate that species-specific traits still modulate realized connectivity. For example, *G. hofrichteri* exhibits slightly weaker population differentiation and a broader distribution range compared to *G. orientalis*, which may reflect differences in dispersal ability or larval ecology. Indeed, *G. hofrichteri* has a wide distribution range and also occurs within the Adriatic Sea, suggesting overall higher dispersal abilities (Wagner et al., 2019, 2021). This result fits observations in the sister genus *Lepadogaster* (Conway et al., 2019), which are also capable of long range dispersal via local oceanic features (Klein et al., 2016). However, within the same genus, differences in breeding season (and concomitant with that larval duration) between sister-species highlight the role of species-specific life-history traits in shaping realized connectivity patterns (Henriques et al., 2002; Tojeira et al., 2012; Wagner et al., 2017).

Together, our results indicate that while oceanographic structure generates highly predictable patterns of connectivity across species, the strength and fine-scale expression of these patterns remain modulated by species-specific biological traits, raising the question of whether such shared connectivity patterns also translate into parallel genomic responses.

### Significant but limited signals of parallel genomic local population divergence

Similar patterns of population structure were associated with a detectable excess of parallel genomic divergence. By focusing on Kythira as a shared focal population across both species, we minimized environmental variation and compared genomic responses under broadly similar oceanographic and ecological conditions. Although most candidate regions were species-specific, overlap among both genomic windows and genes exceeded random expectations, indicating that comparisons involving the same geographic population (Kythira) can leave repeatable genomic signatures across independently evolving lineages.

Notably, one of the overlapping candidate genes was *tuba*, which encodes a core component of the microtubule cytoskeleton. Microtubules play fundamental roles in cell polarity, morphology and tissue organization (Larson et al., 2010), suggesting that some aspects of local divergence may involve shared cellular organizational processes across species. However, the limited number of overlapping genes and the absence of strong functional enrichment prevent more specific biological inference.

Despite this significant overlap, most candidate regions remained species-specific. Candidate regions were distributed across the genome and were predominantly located in non-coding regions, suggesting that local divergence is polygenic and likely mediated largely through regulatory variation rather than a small number of major-effect loci. This interpretation is consistent with the broad and largely unspecific functional enrichment patterns observed, as well as the limited overlap of candidate regions between species. Similar patterns have also been observed in our recent study of convergent adaptation across *Gouania*, where clear phenotypic convergence was accompanied by weak and diffuse genomic signals, likely reflecting the combined effects of polygenic architectures, recombination and evolutionary divergence among lineages (Wagner et al., 2026).

Differences in the extent of divergence between species further suggest an important role for demographic history. *Gouania orientalis* exhibited stronger population structure, higher genome-wide π, and substantially more nearly fixed variants than *G. hofrichteri*. Moreover, nearly fixed variants in *G. orientalis* were frequently clustered within genomic regions, whereas those in *G. hofrichteri* were fewer and more dispersed. Together with differences in genome-wide Tajima’s D, these patterns indicate contrasting demographic histories that likely influenced the accumulation of genomic differentiation, consistent with the expected effects of genetic drift on allele frequency dynamics (Kimura, 1968).

Subtle ecological differences may also contribute to the predominance of species-specific candidate regions. Although both species occur in sympatry and experience broadly similar environmental conditions, subtle differences in microhabitat use could expose populations to distinct selective pressures. Comparable small-scale habitat differentiation has been documented among Adriatic *Gouania* species (Wagner et al., 2023). However, the limited functional overlap between species suggests that similar ecological challenges may often be mediated through different genetic pathways.

Together, these results indicate that similar environmental and oceanographic contexts can generate repeatable genomic responses across independently evolving lineages, but that this predictability is limited. Most genomic divergence remains species-specific and appears to reflect a combination of polygenic architecture, demographic history and lineage-specific ecological responses.

### Species-specific demographic histories contrast with parallel contemporary connectivity

In contrast to the strong concordance observed for contemporary connectivity and aspects of genomic divergence, basin-wide mitochondrial phylogeographic patterns and species-level demographic histories were considerably more heterogeneous. While the COI data support a broad contrast between relatively shallow western/Adriatic structure and deeper eastern Mediterranean structure, they do not show a simple one-to-one correspondence between sympatric species in the placement of individual populations. These mitochondrial patterns and the MSMC trajectories therefore provide broad historical context for our genomic analyses, but also highlight that single-locus phylogeographic structure does not necessarily mirror genome-wide connectivity at fine spatial scales.

Interestingly, the observed patterns closely align with basin-specific palaeoenvironmental histories and are most parsimoniously explained by Quaternary sea-level oscillations, with the Adriatic Sea providing a particularly clear example. During glacial lowstands (ca. 15,000 - 20,000 years ago), the sea level was around 120 m lower than today and large parts of the northern Adriatic were exposed (Lambeck et al., 2014). This drastically reduced the available shallow coastal habitats, leading to population contractions. Consistent with this, *G. adriatica* exhibits a pronounced star-like haplotype network indicative of recent population growth, matching the re-establishment of the modern Adriatic coastline after the last glacial maximum. In contrast, *G. pigra* shows a stronger north–south phylogeographic structure, suggesting either more persistent refugial structure within the basin or a more complex recolonization history. Similar postglacial (northern) expansion dynamics have been reported for other benthic taxa in the Adriatic Sea, indicating that these patterns likely reflect shared responses to basin-specific habitat dynamics rather than species-specific processes alone (Koblmüller et al., 2015; Ledoux et al., 2018; Sefc et al., 2020).

The eastern Mediterranean species further illustrate the limits of phylogeographic predictability. Both *G. orientalis* and *G. hofrichteri* exhibit elevated haplotype diversity and deeper mitochondrial structure, consistent with greater long-term habitat stability in the eastern basin. However, the placement of the populations of Crete and Kythira differed substantially between species, demonstrating that even closely related taxa occupying similar environments can retain distinct historical signatures. Beyond these basin-scale effects, intraspecific structure also aligns with major Mediterranean ecoregions (Spalding et al., 2007) and oceanographic fronts (El-Geziry & Bryden, 2010), further supporting the role of persistent seascape features in shaping connectivity. For example, the clear break between populations from Messina and the French coastline in *G. willdenowi* is consistent with restricted dispersal across major Mediterranean ecogeographic regions, namely the Tyrrhenian and Algero-Provencal basins (Spalding et al., 2007).

Together, the basin-wide analyses reveal a striking contrast with the contemporary genomic results around Crete. Whereas oceanographic conditions appear capable of generating repeatable patterns of connectivity and even partially parallel genomic divergence across species, mitochondrial phylogeographic structure and long-term demographic trajectories are substantially less predictable. These deeper historical signatures likely reflect species-specific responses to Pleistocene environmental change, demographic stochasticity and the persistence of historical population structure. Consequently, predictability in Gouania appears strongest for contemporary processes and progressively weaker when viewed across longer evolutionary timescales.

## CONCLUSION

Our study demonstrates that the predictability of marine population structure varies across evolutionary timescales. At contemporary spatial scales, shared seascape features and oceanographic processes generate remarkably concordant patterns of connectivity across sympatric *Gouania* species. Moreover, shared population contrasts were associated with a significant excess of parallel genomic divergence, indicating that common environmental and demographic contexts can leave partially repeatable signatures across independently evolving lineages. In contrast, mitochondrial phylogeographic patterns and species-level demographic histories were considerably more heterogeneous, highlighting the increasing influence of species-specific historical processes over longer timescales.

For conservation management, these findings emphasize that efficient planning requires integrating oceanographic predictions with species-specific biological information and empirical genetic estimates of realized connectivity (Jahnke et al., 2017; Paterno et al., 2017). Our results further show that genomic data are essential for assessing the evolutionary and adaptive potential of populations at local scales, particularly in rapidly changing environments such as the Mediterranean Sea (Boulanger et al., 2022). This is especially relevant for highly specialized species such as *Gouania*, whose gravel beach habitats are increasingly threatened by beach nourishment and coastal development (Carević, 2020; Carević et al., 2022; Wagner et al., 2023). Accordingly, understanding patterns of connectivity and local genomic divergence will be critical for predicting demographic resilience and preserving evolutionary potential in this endemic Mediterranean genus.

## ACKNOWLEDGMENTS/FUNDING

The authors thank NOBIS-Austria for the financial support (NOBIS-Scholarship to M.W.). M.W. has also been supported with a DOC-Scholarship by the Austrian Academy of Science during his PhD and a YUFE4Postdoc-Scholarship, co-funded by the EU under Marie Sklodowska-Curie Grant Agreement No. 101081327. H.S. acknowledges support from the Research Fund of the University of Antwerp”.

## Notes

### Competing Interest Statement

The authors have declared no competing interest.

## REFERENCES

Agiadi, K., Hohmann, N., Gliozzi, E., Thivaiou, D., Bosellini, F. R., Taviani, M., Bianucci, G., Collareta, A., Londeix, L., Faranda, C., Bulian, F., Koskeridou, E., Lozar, F., Mancini, A. M., Dominici, S., Moissette, P., Campos, I. B., Borghi, E., Iliopoulos, G.,…García-Castellanos, D. (2024). The marine biodiversity impact of the Late Miocene Mediterranean salinity crisis. Science, 385(6712), 986–991. 10.1126/science.adp3703

Alexander, D. H., Novembre, J., & Lange, K. (2009). Fast model-based estimation of ancestry in unrelated individuals. Genome Research, 19(9), 1655–1664. 10.1101/gr.094052.109

Arranz, V., Thakur, V., & Lavery, S. D. (2021). Demographic history, not larval dispersal potential, explains differences in population structure of two New Zealand intertidal species. Marine Biology, 168(7), 105. 10.1007/s00227-021-03891-2

Benjamini, Y., & Hochberg, Y. (1995). Controlling the False Discovery Rate: A Practical and Powerful Approach to Multiple Testing. Journal of the Royal Statistical Society: Series B (Methodological), 57(1), 289–300. 10.1111/j.2517-6161.1995.tb02031.x

Bianchi, C. N., Morri, C., Chiantore, M., Montefalcone, M., Parravicini, V., & Rovere, a. (2012). Mediterranean Sea Biodiversity Between the Legacy From the Past and a Future of Change.

Boulanger, E., Benestan, L., Guerin, P., Dalongeville, A., Mouillot, D., & Manel, S. (2022). Climate differently influences the genomic patterns of two sympatric marine fish species. Journal of Animal Ecology, 91(6), 1180–1195. 10.1111/1365-2656.13623

Brandl, S. J., Goatley, C. H. R., Bellwood, D. R., & Tornabene, L. (2018). The hidden half: Ecology and evolution of cryptobenthic fishes on coral reefs. Biological Reviews, 93(4), 1846–1873. 10.1111/brv.12423

Briggs, J. C. (1955). A monograph of the clingfishes (Order Xenopterygii) (6th ed.). Stanford Ichthyological Bulletin.

Buchfink, B., Reuter, K., & Drost, H.-G. (2021). Sensitive protein alignments at tree-of-life scale using DIAMOND. Nature Methods, 18(4), 366–368. 10.1038/s41592-021-01101-x

Carević, D. (2020). Održiva gradnja nasutih plaža Beachex 2019-2023. Građevinar, 72(12), 1173–1179.

Carević, D., Bogovac, T., Bujak, D., & Ilić, S. (2022). Analiza stanja dohranjivanja i nasipavanja plaža u Hrvatskoj. Građevinar, 74(1), 71–76.

Cingolani, P., Platts, A., Wang, L. L., Coon, M., Nguyen, T., Wang, L., Land, S. J., Lu, X., & Ruden, D. M. (2012). A program for annotating and predicting the effects of single nucleotide polymorphisms, SnpEff: SNPs in the genome of *Drosophila melanogaster* strain w1118; iso-2; iso-3. Fly, 6(2), 80–92. 10.4161/fly.19695

Clement, M., Snell, Q., Walke, P., Posada, D., & Crandall, K. (2002). TCS: Estimating gene genealogies. *Proceedings - International Parallel and Distributed Processing Symposium*, IPDPS 2002, 184. 10.1109/IPDPS.2002.1016585

Clementi, E., Drudi, M., Aydogdu, A., Moulin, A., Grandi, A., Mariani, A., Goglio, A. C., Pistoia, J., Miraglio, P., Lecci, R., Palermo, F., Coppini, G., Masina, S., & Pinardi, N. (2023). *Mediterranean Sea Physical Analysis and Forecast (CMEMS MED-Physics, EAS8 system): MEDSEA_ANALYSISFORECAST_PHY_006_013* (Version 1) [Dataset]. Copernicus Marine Service (CMS). 10.25423/CMCC/MEDSEA_ANALYSISFORECAST_PHY_006_013_EAS8

Conway, K. W., Moore, G. I., & Summers, A. P. (2019). A new genus and two new species of miniature clingfishes from temperate southern Australia (teleostei, gobiesocidae). ZooKeys, 2019(864), 35–65. 10.3897/zookeys.864.34521

Cowen, R. K., Gawarkiewicz, G., Pineda, J., Thorrold, S. R., & Werner, F. E. (2007). Population Connectivity in Marine Systems: An Overview. Oceanography, 20(3), 14–21. 10.5670/oceanog.2009.01

Danecek, P., Auton, A., Abecasis, G., Albers, C. A., Banks, E., DePristo, M. A., Handsaker, R. E., Lunter, G., Marth, G. T., Sherry, S. T., McVean, G., Durbin, R., & 1000 Genomes Project Analysis Group. (2011). The variant call format and VCFtools. Bioinformatics, 27(15), 2156–2158. 10.1093/bioinformatics/btr330

Danecek, P., Bonfield, J. K., Liddle, J., Marshall, J., Ohan, V., Pollard, M. O., Whitwham, A., Keane, T., McCarthy, S. A., Davies, R. M., & Li, H. (2021). Twelve years of SAMtools and BCFtools. GigaScience, 10(2), giab008. 10.1093/gigascience/giab008

Dawson, M. N. (2012). Parallel phylogeographic structure in ecologically similar sympatric sister taxa. Molecular Ecology, 21(4), 987–1004. 10.1111/j.1365-294X.2011.05417.x

Delandmeter, P., & Van Sebille, E. (2019). The Parcels v2.0 Lagrangian framework: New field interpolation schemes. Geoscientific Model Development, 12(8), 3571–3584. 10.5194/gmd-12-3571-2019

Deli, T., Kiel, C., & Schubart, C. D. (2019). Phylogeographic and evolutionary history analyses of the warty crab Eriphia verrucosa (Decapoda, Brachyura, Eriphiidae) unveil genetic imprints of a late Pleistocene vicariant event across the Gibraltar Strait, erased by postglacial expansion and admixture among refugial lineages. BMC Evolutionary Biology, 19(1), 105. 10.1186/s12862-019-1423-2

Edgar, R. C. (2004). MUSCLE: multiple sequence alignment with high accuracy and high throughput. Nucleic Acids Research, 32(5), 1792–1797. 10.1093/chemse/bjs128

El-Geziry, T. M., & Bryden, I. G. (2010). The circulation pattern in the Mediterranean Sea: Issues for modeller consideration. Journal of Operational Oceanography, 3(2), 39–46. 10.1080/1755876X.2010.11020116

Giakoumi, S., Sini, M., Gerovasileiou, V., Mazor, T., Beher, J., Possingham, H. P., Abdulla, A., Çinar, M. E., Dendrinos, P., Gucu, A. C., Karamanlidis, A. A., Rodic, P., Panayotidis, P., Taskin, E., Jaklin, A., Voultsiadou, E., Webster, C., Zenetos, A., & Katsanevakis, S. (2013). Ecoregion-Based Conservation Planning in the Mediterranean: Dealing with Large-Scale Heterogeneity. PLoS ONE, 8(10), e76449. 10.1371/journal.pone.0076449

Hellberg, M. E. (2009). Gene Flow and Isolation among Populations of Marine Animals. Annual Review of Ecology, Evolution, and Systematics, 40(1), 291–310. 10.1146/annurev.ecolsys.110308.120223

Henriques, M., Lourenço, R., Almada, F., Calado, G., Gonçalves, D., Guillemaud, T., Cancela, M. L., & Almada, V. C. (2002). A revision of the status of Lepadogaster lepadogaster (Teleostei: Gobiesocidae): sympatric subspecies or a long misunderstood blend of species? Biological Journal of the Linnean Society, 76(3), 327–338. 10.1111/j.1095-8312.2002.tb01700.x

Hudson, R. R., Boos, D. D., & Kaplan, N. L. (1992). A statistical test for detecting geographic subdivision. Molecular Biology and Evolution, 9(1), 138–151. 10.1093/oxfordjournals.molbev.a040703

Jahnke, M., Casagrandi, R., Melià, P., Schiavina, M., Schultz, S. T., Zane, L., & Procaccini, G. (2017). Potential and realized connectivity of the seagrass *Posidonia oceanica* and their implication for conservation. Diversity and Distributions, 23(12), 1423–1434. 10.1111/ddi.12633

Kato, S., Arakaki, S., Nagano, A. J., Kikuchi, K., & Hirase, S. (2025). Discordant Phylogeographic Patterns in Ecologically Similar Sympatric Sister Species: Revisiting the Null Hypothesis of Comparative Phylogeography. Journal of Biogeography, 52(12), e70112. 10.1111/jbi.70112

Kimura, M. (1968). Evolutionary Rate at the Molecular Level. Nature, 217(5129), 624–626. 10.1038/217624a0

Klein, M., Teixeira, S., Assis, J., Serrão, E. A., Gonçalves, E. J., & Borges, R. (2016). High Interannual Variability in Connectivity and Genetic Pool of a Temperate Clingfish Matches Oceanographic Transport Predictions. Plos One, 11(12), e0165881. 10.1371/journal.pone.0165881

Koblmüller, S., Steinwender, B., Weiß, S., & Sefc, K. M. (2015). Gene flow, population growth and a novel substitution rate estimate in a subtidal rock specialist, the black-faced blenny *Tripterygion delaisi* (Perciformes, Blennioidei, Tripterygiidae) from the Adriatic Sea. Journal of Zoological Systematics and Evolutionary Research, 53(4), 291–299. 10.1111/jzs.12110

Kousteni, V., Kasapidis, P., Kotoulas, G., & Megalofonou, P. (2015). Strong population genetic structure and contrasting demographic histories for the small-spotted catshark (Scyliorhinus canicula) in the Mediterranean Sea. Heredity, 114(3), 333–343. 10.1038/hdy.2014.107

Kumar, S., Stecher, G., & Tamura, K. (2016). MEGA7: Molecular Evolutionary Genetics Analysis Version 7.0 for Bigger Datasets. Molecular Biology and Evolution, 33(7), 1870–1874. 10.1093/molbev/msw054

Lambeck, K., Rouby, H., Purcell, A., Sun, Y., & Sambridge, M. (2014). Sea level and global ice volumes from the Last Glacial Maximum to the Holocene. Proceedings of the National Academy of Sciences, 111(43), 15296–15303. 10.1073/pnas.1411762111

Lange, M., & Van Sebille, E. (2017). Parcels v0.9: Prototyping a Lagrangian ocean analysis framework for the petascale age. Geoscientific Model Development, 10(11), 4175–4186. 10.5194/gmd-10-4175-2017

Larson, T. A., Gordon, T. N., Lau, H. E., & Parichy, D. M. (2010). Defective adult oligodendrocyte and Schwann cell development, pigment pattern, and craniofacial morphology in puma mutant zebrafish having an alpha tubulin mutation. Developmental Biology, 346(2), 296–309. 10.1016/j.ydbio.2010.07.035

Ledoux, J., Frleta-Valić, M., Kipson, S., Antunes, A., Cebrian, E., Linares, C., Sánchez, P., Leblois, R., & Garrabou, J. (2018). Postglacial range expansion shaped the spatial genetic structure in a marine habitat-forming species: Implications for conservation plans in the Eastern Adriatic Sea. Journal of Biogeography, 45(12), 2645–2657. 10.1111/jbi.13461

Leigh, J. W., & Bryant, D. (2015). POPART: Full-feature software for haplotype network construction. Methods in Ecology and Evolution, 6(9), 1110–1116. 10.1111/2041-210X.12410

Lescroart, J., Bonilla-Sánchez, A., Napolitano, C., Buitrago-Torres, D. L., Ramírez-Chaves, H. E., Pulido-Santacruz, P., Murphy, W. J., Svardal, H., & Eizirik, E. (2023). Extensive Phylogenomic Discordance and the Complex Evolutionary History of the Neotropical Cat Genus *Leopardus*. Molecular Biology and Evolution, 40(12), msad255. 10.1093/molbev/msad255

Macpherson, E., & Raventos, N. (2006). Relationship between pelagic larval duration and geographic distribution of Mediterranean littoral fishes. Marine Ecology Progress Series, 327, 257–265.

Malinsky, M., Matschiner, M., & Svardal, H. (2021). Dsuite—Fast D-statistics and related admixture evidence from VCF files. Molecular Ecology Resources, 21(2), 584–595. 10.1111/1755-0998.13265

Malinsky, M., Svardal, H., Tyers, A. M., Miska, E. A., Genner, M. J., Turner, G. F., & Durbin, R. (2018). Whole-genome sequences of Malawi cichlids reveal multiple radiations interconnected by gene flow. Nature Ecology & Evolution, 2(12), 1940–1955. 10.1038/s41559-018-0717-x

Mölder, F., Jablonski, K. P., Letcher, B., Hall, M. B., Tomkins-Tinch, C. H., Sochat, V., Forster, J., Lee, S., Twardziok, S. O., Kanitz, A., Wilm, A., Holtgrewe, M., Rahmann, S., Nahnsen, S., & Köster, J. (2021). Sustainable data analysis with Snakemake. F1000Research, 10, 33. 10.12688/f1000research.29032.1

Ollé-Vilanova, J., Pérez-Bielsa, N., Araguas, R. M., Sanz, N., Saber, S., Macías, D., & Viñas, J. (2022). Larval Retention and Homing Behaviour Shape the Genetic Structure of the Bullet Tuna (Auxis rochei) in the Mediterranean Sea. Fishes, 7(5), 300. 10.3390/fishes7050300

Palumbi, S. R. (2003). POPULATION GENETICS, DEMOGRAPHIC CONNECTIVITY, AND THE DESIGN OF MARINE RESERVES. Ecological Applications, 13(sp1), 146–158. 10.1890/1051-0761(2003)013%5B0146:PGDCAT%5D2.0.CO;2

Pascual, M., Rives, B., Schunter, C., & Macpherson, E. (2017). Impact of life history traits on gene flow: A multispecies systematic review across oceanographic barriers in the Mediterranean Sea. PLOS ONE, 12(5), e0176419. 10.1371/journal.pone.0176419

Paterno, M., Schiavina, M., Aglieri, G., Ben Souissi, J., Boscari, E., Casagrandi, R., Chassanite, A., Chiantore, M., Congiu, L., Guarnieri, G., Kruschel, C., Macic, V., Marino, I. A. M., Papetti, C., Patarnello, T., Zane, L., & Melià, P. (2017). Population genomics meet Lagrangian simulations: Oceanographic patterns and long larval duration ensure connectivity among Paracentrotus lividus populations in the Adriatic and Ionian seas. Ecology and Evolution, 7(8), 2463–2479. 10.1002/ece3.2844

Poulin, P.-M., Bussani, A., Gerin, R., Jungwirth, R., Mauri, E., Menna, M., & Notarstefano, G. (2013). Mediterranean Surface Currents Measured with Drifters: From Basin to Subinertial Scales. Oceanography, 26(1), 38–47. 10.5670/oceanog.2013.03

Purcell, S., Neale, B., Todd-Brown, K., Thomas, L., Ferreira, M. A. R., Bender, D., Maller, J., Sklar, P., de Bakker, P. I. W., Daly, M. J., & Sham, P. C. (2007). PLINK: A Tool Set for Whole-Genome Association and Population-Based Linkage Analyses. The American Journal of Human Genetics, 81(3), 559–575. 10.1086/519795

Reimand, J., Kull, M., Peterson, H., Hansen, J., & Vilo, J. (2007). G:Profiler-a web-based toolset for functional profiling of gene lists from large-scale experiments. Nucleic Acids Research, 35(SUPPL.2). 10.1093/nar/gkm226

Rhie, A., McCarthy, S. A., Fedrigo, O., Damas, J., Formenti, G., Koren, S., Uliano-Silva, M., Chow, W., Fungtammasan, A., Kim, J., Lee, C., Ko, B. J., Chaisson, M., Gedman, G. L., Cantin, L. J., Thibaud-Nissen, F., Haggerty, L., Bista, I., Smith, M.,…Jarvis, E. D. (2021). Towards complete and error-free genome assemblies of all vertebrate species. Nature, 592(7856), 737–746. 10.1038/s41586-021-03451-0

Schiffels, S., & Wang, K. (2020). MSMC and MSMC2: The multiple sequentially Markovian coalescent. In Methods in Molecular Biology (pp. 147–166). 10.1007/978-1-0716-0199-0_7

Schroeder, K., Garcìa-Lafuente, J., Josey, S. A., Artale, V., Nardelli, B. B., Carrillo, A., Gačić, M., Gasparini, G. P., Herrmann, M., Lionello, P., Ludwig, W., Millot, C., Özsoy, E., Pisacane, G., Sánchez-Garrido, J. C., Sannino, G., Santoleri, R., Somot, S., Struglia, M.,…Zodiatis, G. (2012). Circulation of the Mediterranean Sea and its Variability. In The Climate of the Mediterranean Region (pp. 187–256). Elsevier. 10.1016/B978-0-12-416042-2.00003-3

Sefc, K. M., Wagner, M., Zangl, L., Weiß, S., Steinwender, B., Arminger, P., Weinmaier, T., Balkic, N., Kohler, T., & Inthal, S. (2020). Phylogeographic structure and population connectivity of a small benthic fish (Tripterygion tripteronotum) in the Adriatic Sea. Journal of Biogeography, 47(11), 2502–2517.

Selkoe, K. A., Henzler, C. M., & Gaines, S. D. (2008). Seascape genetics and the spatial ecology of marine populations. Fish and Fisheries, 9(4), 363–377. 10.1111/j.1467-2979.2008.00300.x

Spalding, M. D., Fox, H. E., Allen, G. R., Davidson, N., Ferdaña, Z. A., Finlayson, M., Halpern, B. S., Jorge, M. A., Lombana, A., Lourie, S. A., Martin, K. D., McManus, E., Molnar, J., Recchia, C. A., & Robertson, J. (2007). Marine Ecoregions of the World: A Bioregionalization of Coastal and Shelf Areas. BioScience, 57(7), 573–583. 10.1641/B570707

Tajima, F. (1989). Statistical method for testing the neutral mutation hypothesis by DNA polymorphism. Genetics, 123(3), 585–595. 10.1093/genetics/123.3.585

Tojeira, I., Faria, A. M., Henriques, S., Faria, C., & Gonçalves, E. J. (2012). Early development and larval behaviour of two clingfishes, Lepadogaster purpurea and Lepadogaster lepadogaster (Pisces: Gobiesocidae). Environmental Biology of Fishes, 93(4), 449–459. 10.1007/s10641-011-9935-7

Wagner, M., Benac, Č., Pamić, M., Bračun, S., Ladner, M., Plakolm, P., Koblmüller, S., Svardal, H., & Brandl, S. (2023). Microhabitat partitioning between sympatric intertidal fish species highlights the importance of sediment composition in gravel beach conservation. Ecology and Evolution, 13, e10302.

Wagner, M., Bračun, S., Kovačić, M., Iglésias, S. P., Sellos, D. Y., Zogaris, S., & Koblmüller, S. (2017). *Lepadogaster purpurea* (Actinopterygii: Gobiesociformes: Gobiesocidae) from the eastern mediterranean sea: Significantly extended distribution range. Acta Ichthyologica et Piscatoria, 47(4), 417–421. 10.3750/AIEP/02244

Wagner, M., Bračun, S., Skofitsch, G., Kovačić, M., Zogaris, S., Iglésias, S. P., Sefc, K. M., & Koblmüller, S. (2019). Diversification in gravel beaches: A radiation of interstitial clingfish (*Gouania*, Gobiesocidae) in the Mediterranean Sea. Molecular Phylogenetics and Evolution, 139, 106525. 10.1016/j.ympev.2019.106525

Wagner, M., Kovačić, M., & Koblmüller, S. (2021). Unravelling the taxonomy of an interstitial fish radiation: Three new species of *Gouania* (Teleostei: Gobiesocidae) from the Mediterranean Sea and redescriptions of *G. willdenowi* and *G. pigra*. Journal of Fish Biology, 98(1), 64–88. 10.1111/jfb.14558

Wagner, M., Resl, P., Klar, N., Huie, J. M., Bista, I., McCarthy, S., Smith, M., Durbin, R., Koblmüller, S., & Svardal, H. (2026). The genomics of convergent adaptation to intertidal gravel beaches in Mediterranean clingfishes. *Genome Biology and Evolution*, evag031. 10.1093/gbe/evag031

Weersing, K., & Toonen, R. J. (2009). Population genetics, larval dispersal, and connectivity in marine systems. Marine Ecology Progress Series, 393, 1–12. 10.3354/meps08287

Yi, X., Liang, Y., Huerta-Sanchez, E., Jin, X., Cuo, Z. X. P., Pool, J. E., Xu, X., Jiang, H., Vinckenbosch, N., Korneliussen, T. S., Zheng, H., Liu, T., He, W., Li, K., Luo, R., Nie, X., Wu, H., Zhao, M., Cao, H.,…Wang, J. (2010). Sequencing of 50 Human Exomes Reveals Adaptation to High Altitude. Science, 329(5987), 75–78. 10.1126/science.1190371

