## Supplementary Figures for "Parallel connectivity and genomic divergence but heterogeneous demographic history in sympatric Mediterranean clingfishes"

for

Maximilian Wagner et al.

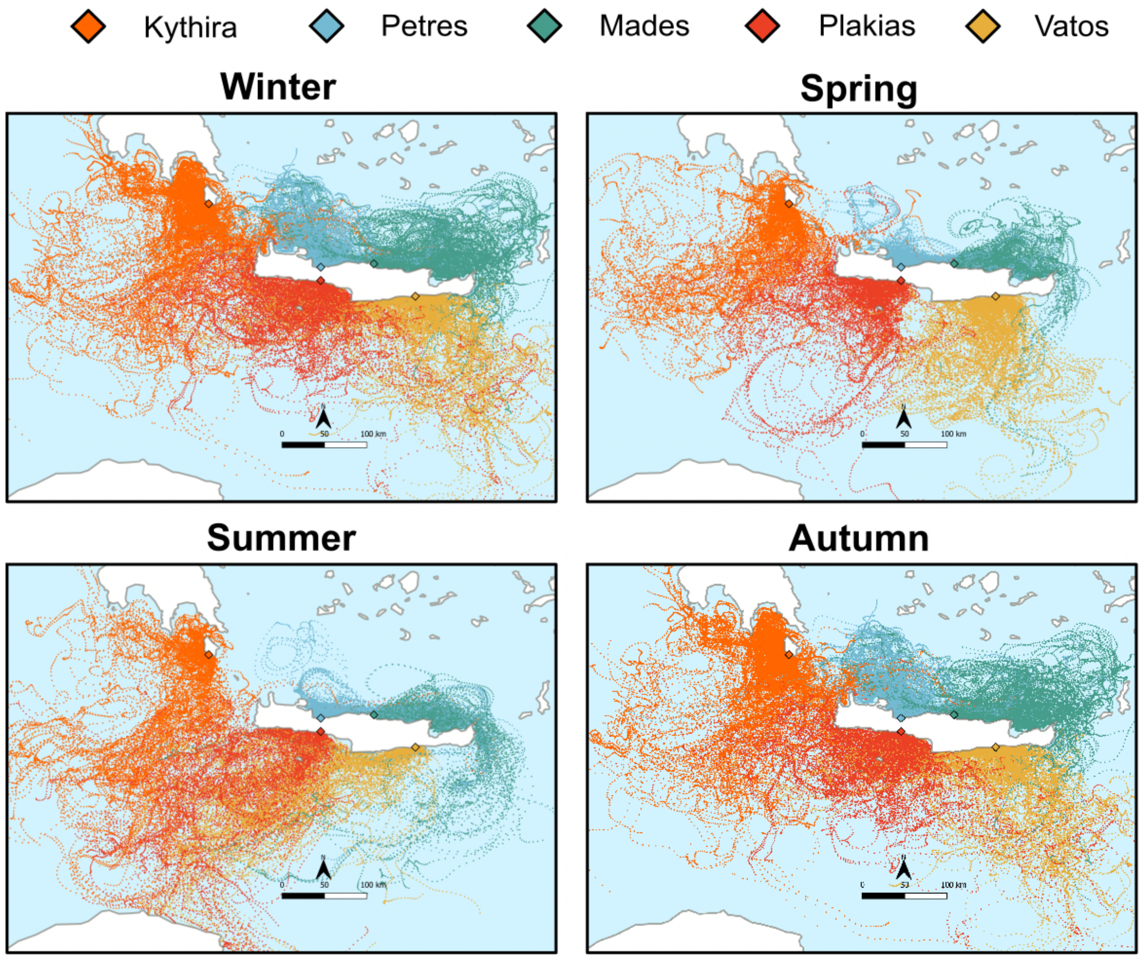


**Figure S1 | Lagrangian simulations of passive larval drift around the island of Crete PLD 26 days**. Propagules were released from the four populations on the island of Crete and the island of Kythira. Each dot represents a time stamp of 6 hours. We conducted four independent analyses spanning over the four seasons: Winter (November – March), spring (March – June), summer (June – September) and autumn (September – Novermber), to test if potential seasonality in the breeding season might explain subtle genetic populations structure differences between sympatric *Gouania orienatlis* and *G. hofrichteri*. Larvae were released every day and drifted for 26 days.

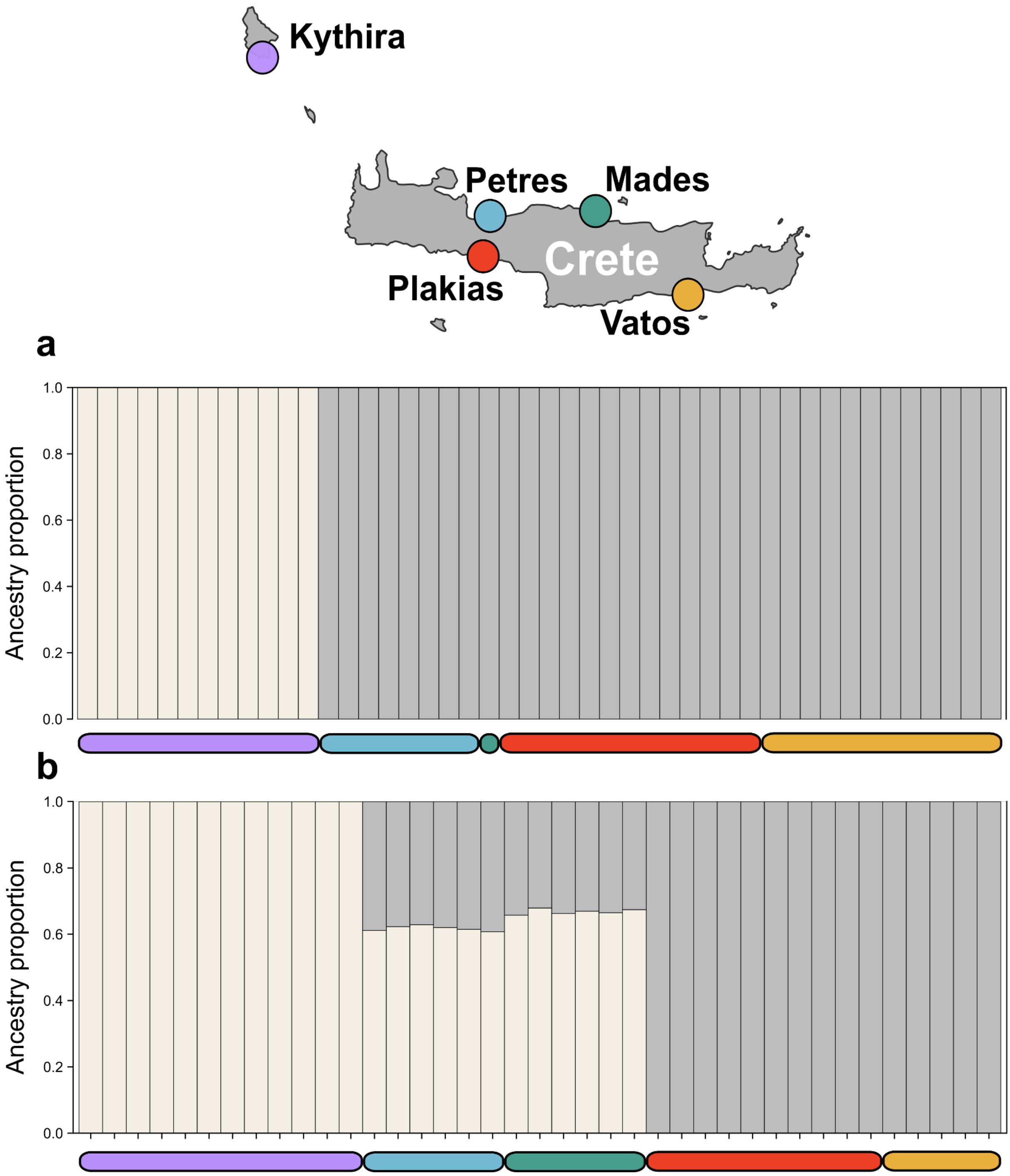


**Figure S2 | Admixture plots show strong geographic structure** for **(a)** *Gouania orientalis*, which separates two main haplotypes, the island of Kythira and the island of Crete. **(b)** *G. hofrichteri* populations from the North of Crete are admixed with the population of Kythira. The colors correspond to different populations in the map above.

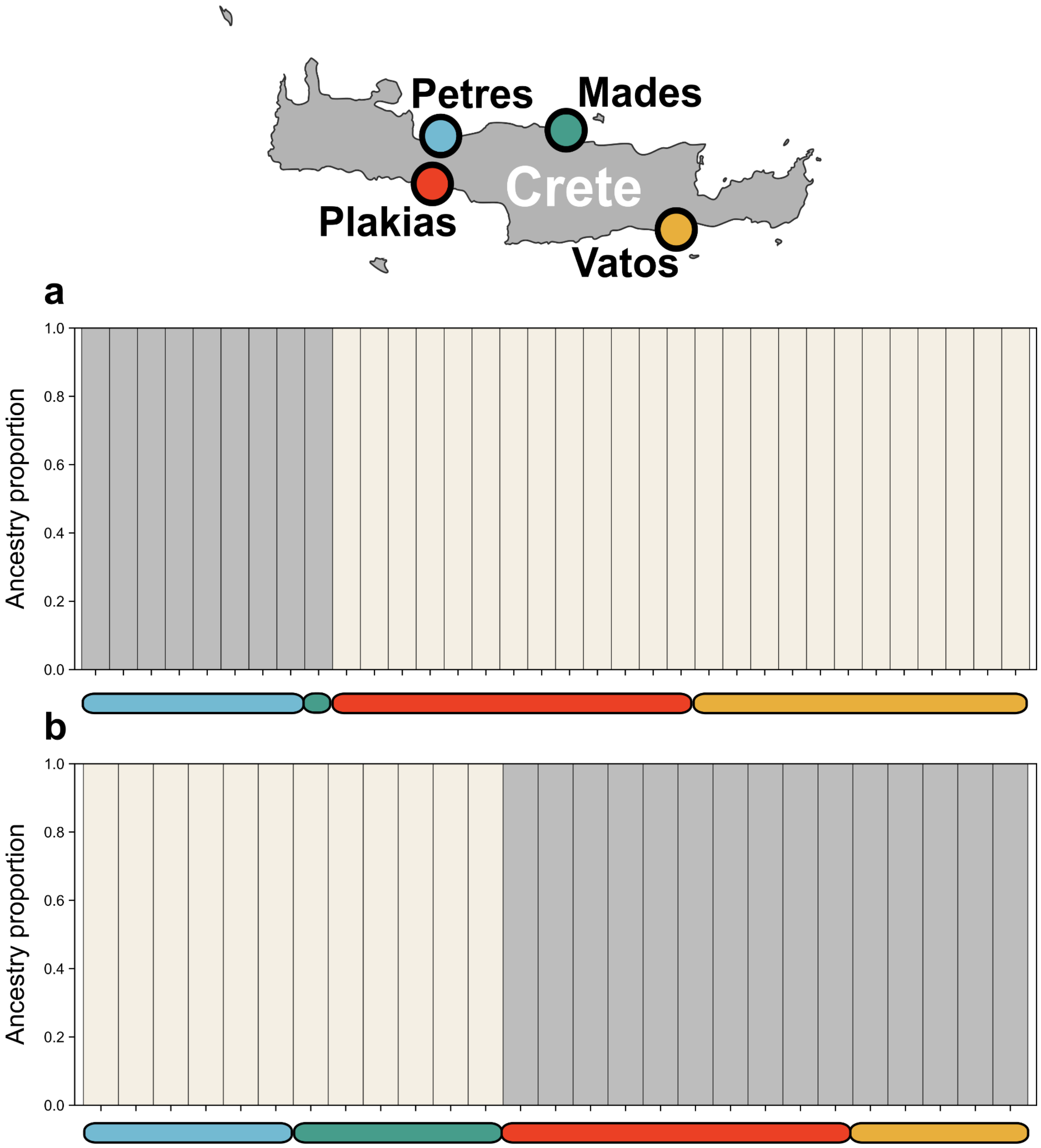


**Figure S3 | Admixture plots based on Crete populations only** reveal strong geographic structure separating two main ancestry components corresponding to northern versus southern Crete in both (a) *Gouania orientalis* and (b) *G. hofrichteri*. In both species, populations from northern Crete show admixture with the Kythira population. Colors correspond to the sampling locations shown in the map above.

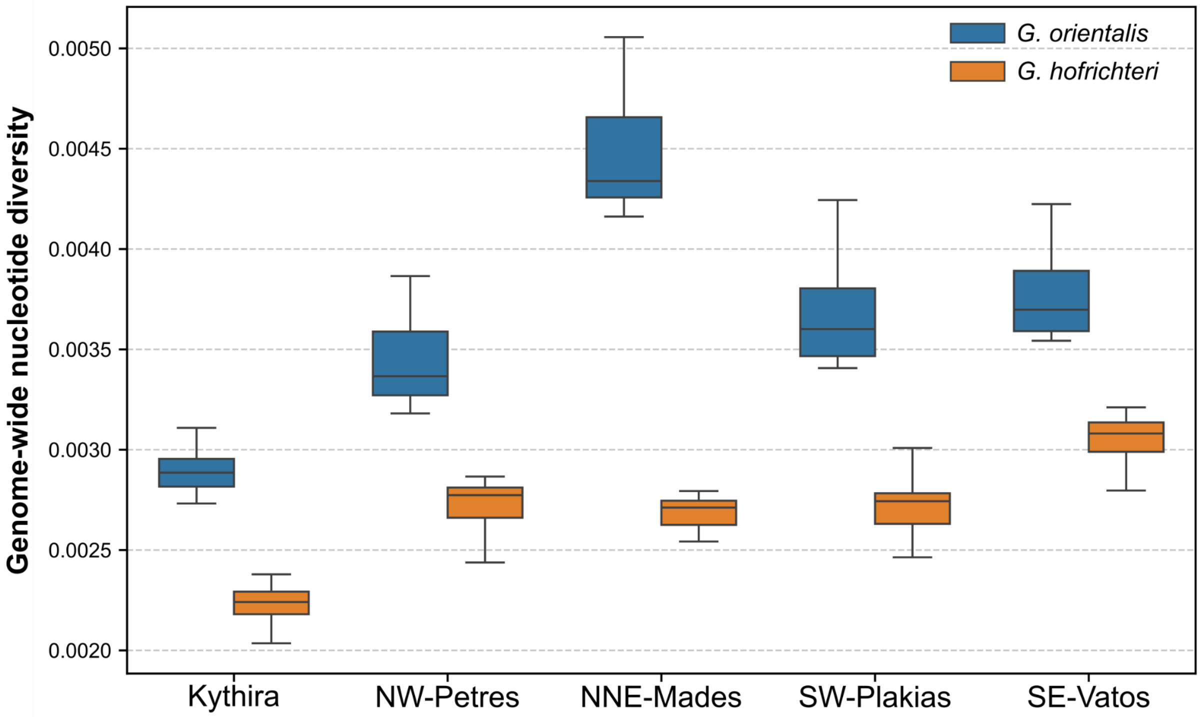


**Figure S4 | Genome-wide nucleotide diversity per population and species.** Values were obtained in 60 kb windows along the genome and averaged according to chromosomes.

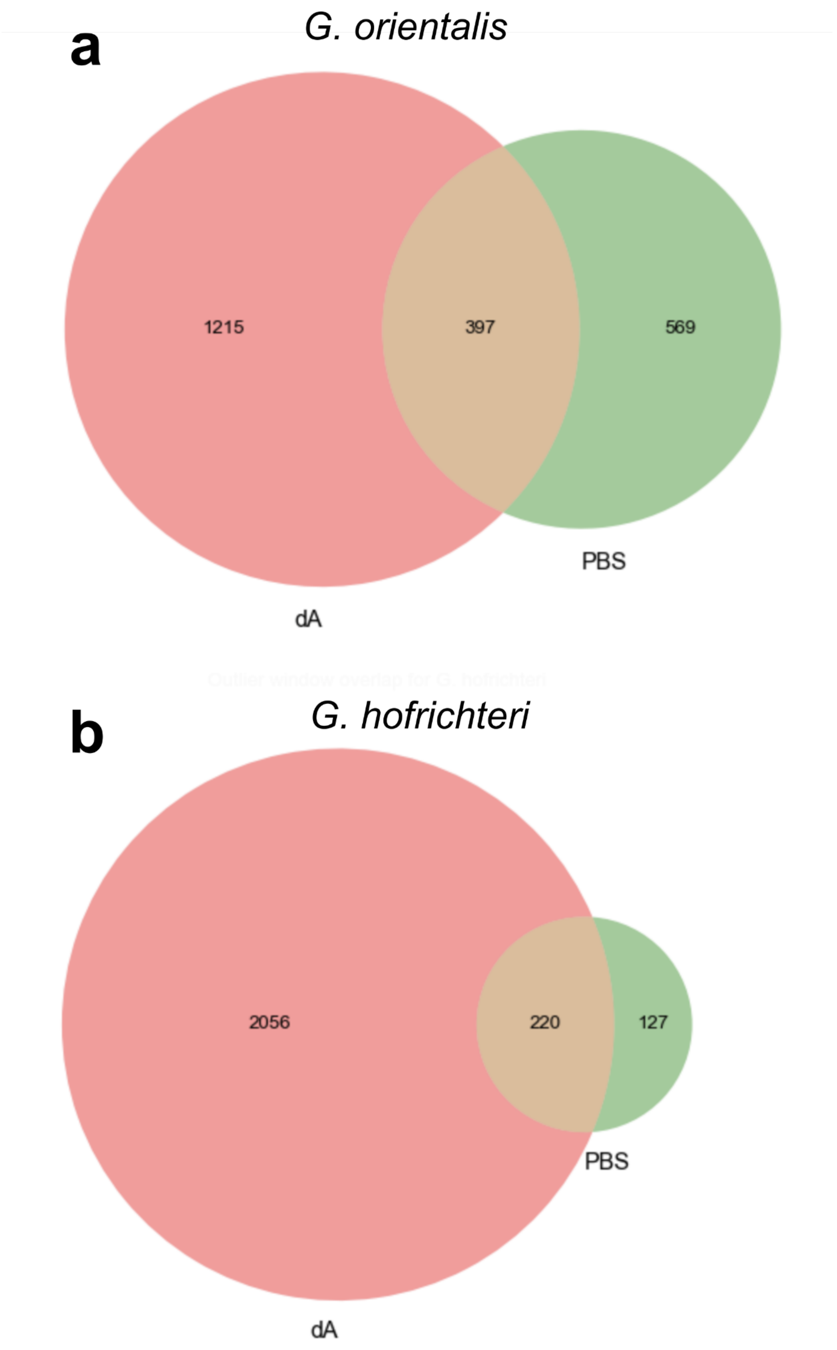


**Figure S5 |** Overlap in the number of outlier $d_A$ and PBS windows for (a) *Gouania orientalis* and (b) *G. hofrichteri*.

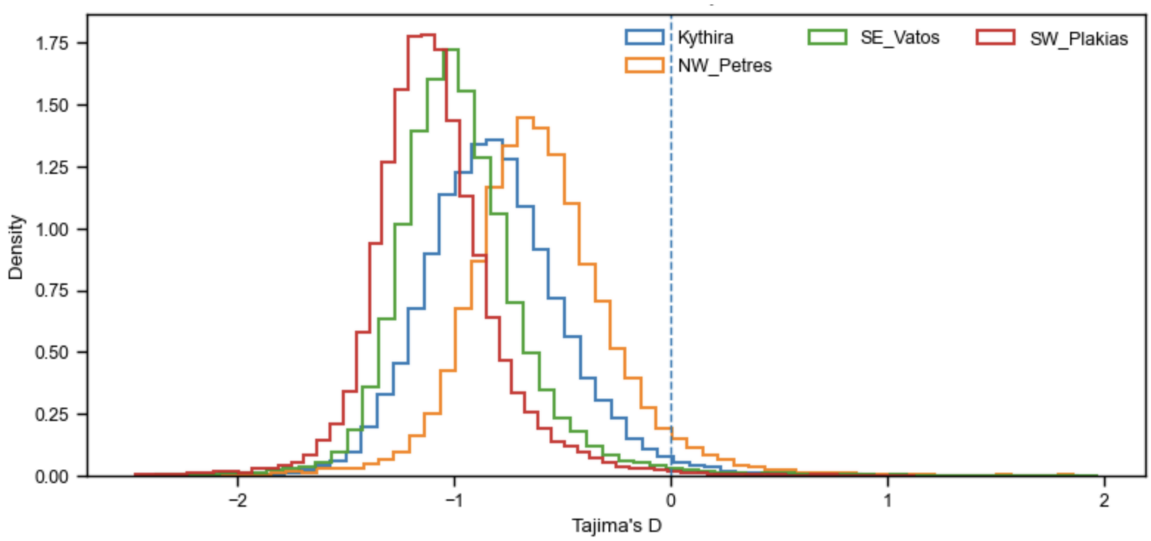


**Figure S6 | Genome-wide distribution of Tajima's D in *Gouania orientalis*.** Density distributions of Tajima's D calculated in sliding windows for the Kythira population and three Cretan populations (Petres, Vatos, and Plakias). Across all populations, Tajima's D values are predominantly negative, indicating an excess of rare alleles consistent with historical demographic expansion or pervasive purifying selection. Differences among populations are modest and largely reflect shifts in central tendency rather than distinct population-specific signals. The dashed vertical line indicates Tajima's D = 0.

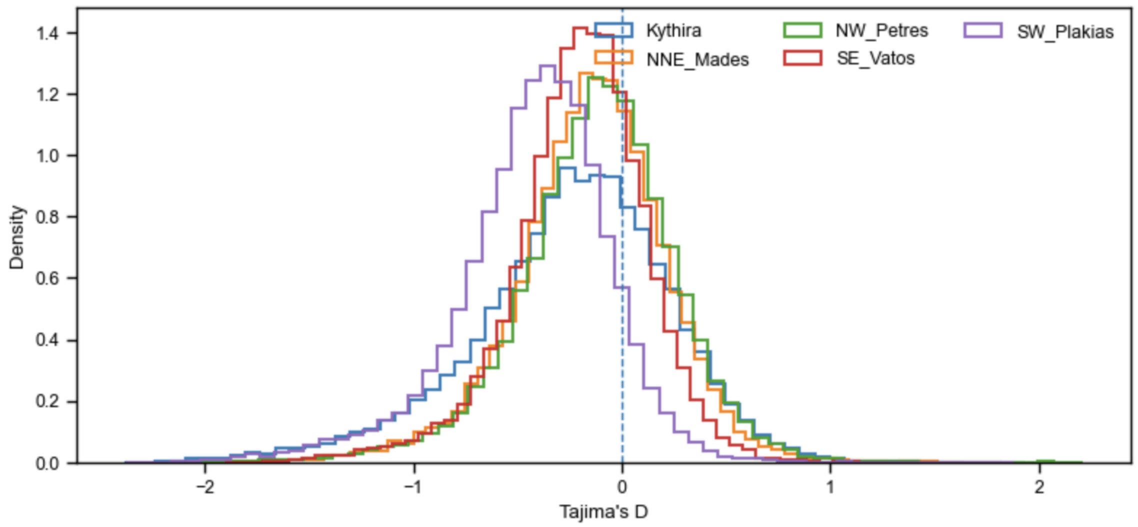


**Figure S6 | Genome-wide distribution of Tajima's D in *Gouania hofrichteri*.**
Density distributions of Tajima's D calculated in sliding windows for the Kythira population and four Cretan populations (Petres, Mades, Vatos, and Plakias). In contrast to *G. orientalis*, Tajima's D values in *G. hofrichteri* are centered closer to zero across populations, consistent with comparatively stable demographic histories or weaker genome-wide signals of recent expansion. Population-level differences are subtle and primarily reflect shifts in distribution width rather than pronounced population-specific deviations. The dashed vertical line indicates Tajima's D = 0.
